# Variations in regulatory sequences rewire gene networks that differentiate wild and cultivated rice for leaf development and photosynthesis

**DOI:** 10.1101/2025.08.07.669153

**Authors:** Vikram Jathar, Mahesh Kumar Panda, AT Vivek, Rajarshi Sanyal, Anurag V Daware, Aditi Dwivedi, Ruchi Rani, Shailesh Kumar, Aashish Ranjan

**Affiliations:** BRIC-National Institute of Plant Genome Research, Aruna Asaf Ali Marg, New Delhi 110067, India; Institute of Plant Breeding, Seed Science and Population Genetics, University of Hohenheim, Stuttgart 70599, Germany

**Keywords:** Leaf development, Photosynthesis, Cultivated and wild rice, Gene regulatory networks, Transcription factors, Regulatory sequence

## Abstract

A comprehensive understanding of gene regulatory networks and the key regulators underlying developmental and physiological progression along the leaf developmental gradient is crucial for optimizing photosynthetic competence. Comparisons of developmental and photosynthetic features across successive leaf developmental stages revealed pronounced differences between wild rice (*Oryza australiensis*) and three cultivated rice accessions. Global gene expression profiling identified three major transcriptional phases across leaf stages: a predominance of developmental genes at SAM+Pi (initiating primordia) and P3; genes for photosynthetic transition at P3 and P4; and enriched core photosynthetic genes at P4 and P5. *O. australiensis* showed a more prominent expression of developmental and photosynthetic genes than cultivated accessions across leaf stages. Multivariate analysis further supported a distinct transcriptional landscape in *O. australiensis* compared with cultivated accessions. Integration of gene expression with species-specific variations in regulatory sequences, derived from synteny-anchored orthology analysis, generated stage-resolved gene regulatory networks and identified key transcription factors (TFs) mediating the developmental and physiological progression. A cross-species comparison of TF–target regulatory networks revealed extensive stage- and accession-dependent rewiring of the regulatory networks of key TFs, with strong regulatory divergence in *O. australiensis*. Gene silencing of two candidate TFs, OsDOF8 and OsARID2, altered the rice photosynthetic competence with accession-specific effects. Taken together, the study highlights promoter-driven regulatory divergence as a potential mechanism underlying developmental and photosynthetic differences among the rice accessions. The resource is available through an interactive public database, Rice DEV-LEAF (https://nipgr.ac.in/DEV-LEAF/), enabling exploration of gene expression dynamics and regulatory networks across the rice leaf developmental gradient.

## Introduction

Leaf developmental attributes strongly influence photosynthetic efficiency (Onoda et al., 2017; Tamang et al., 2023; Aneja et al., 2025). Leaf morphological features, such as width and thickness, and anatomical features, such as mesophyll cell differentiation and vascular development, along with stomatal and chloroplast features, integrate leaf development with photosynthetic competence (Ouyang et al., 2022; Xiao et al., 2023). Variations in leaf developmental features, both within and across species, have been shown to influence photosynthetic efficiency (Xiao et al., 2023; Nikolopoulos et al., 2024; Retta et al., 2024). For example, Kranz anatomy in C_4_ plants offers a distinct advantage in CO_2_ assimilation and overall photosynthetic capacity compared to C_3_-specific anatomy (Gowik and Westhoff, 2011; Acevedo-Siaca et al., 2020). Similarly, targeted modification of key leaf developmental regulators has been shown to alter leaf architecture and improve photosynthetic performance (Wang et al., 2017; Lawrence et al., 2021; Østerlund et al., 2025). Therefore, a comprehensive genetic understanding of developmental attributes across successive leaf growth stages is crucial for optimizing photosynthetic efficiency.

Leaf development follows a coordinated sequence of spatiotemporal events, including cell proliferation, elongation, and differentiation. Initiation of a leaf primordium at the shoot apical meristem (SAM) marks the beginning of leaf development that follows a series of distinct developmental stages, classified as Primordia 0 (P0) to P6, with specific developmental and physiological features (Gong et al., 2024; Wu et al., 2026). The early leaf developmental stages (P0 to P2) are marked by the establishment of axial polarity and the initiation of cellular differentiation, followed by the initiation of the stomatal lineage and the differentiation of vascular tissues and plastids at P3–P4 (van Campen et al., 2016; Perico et al., 2024; Sun et al., 2024; Wu et al., 2026). P5 and P6 mark the completion of leaf growth and the establishment of photosynthetically competent leaves (Wu et al., 2026). While the photosynthetic transition in a leaf happens between the P3 and P4, the early developmental events are critical for the photosynthetic functions of a leaf. These distinct leaf developmental stages provide a structured framework for understanding the physiological transition from leaf initiation to photosynthetic competency (Hill and Lord, 1990).

Each leaf developmental stage is characterized by distinct spatiotemporal gene expression patterns that drive cellular and physiological progression underlying leaf structure and function (van Campen et al., 2016; Miya et al., 2021). Coordinated gene expression patterns across the stages, regulated through both cis- and trans-regulatory elements, are critical for the developmental and physiological progression of a leaf (Singh et al., 2023; Swift et al., 2024). Comparative transcriptomic analyses across different leaf developmental stages have provided valuable insights into gene regulatory networks and the roles of cis-regulatory elements in governing leaf developmental traits (Borba et al., 2023; Singh et al., 2023; Swift et al., 2024). Such comparative developmental analyses of C_3_ and C_4_ grasses have identified key regulators associated with the establishment of C_4_ photosynthesis (Wang et al., 2014). A similar study in rice found the photosynthetic transition between P3 and P4 stages of leaf development (van Campen et al., 2016; Miya et al., 2021). However, integrating the leaf developmental framework with photosynthetic transition and efficiency across contrasting *Oryza* accessions could provide a unified framework for understanding the progression of rice leaves from leaf growth to photosynthetic maturity. Moreover, identifying the gene regulatory networks and key regulators governing the progression would be instrumental in optimizing leaf development and photosynthetic efficiency.

Wild relatives of rice, with remarkably distinct developmental and physiological features compared to cultivated rice varieties, offer a promising resource for comparative studies to dissect the genetic basis of developmental and physiological progression in rice leaves (Kiran et al., 2013; Giuliani et al., 2013; Mathan et al., 2021a; Mathan et al., 2021b; Jathar et al., 2022). While previous studies have identified photosynthetically efficient wild rice accessions and the underlying photochemical and biochemical bases, a comprehensive integration of leaf development and photosynthesis, along with the underlying gene regulatory networks, remains limited across the leaf developmental gradient. In this study, integration of developmental and physiological features with the gene expression profile revealed a clear progression from developmental programs to the establishment of photosynthetic competence along the leaf developmental gradient across selected contrasting cultivated and wild rice accessions. Comparative analyses of Gene Regulatory Networks (GRNs) further revealed both conserved and accession-specific regulatory interactions, with extensive rewiring of transcription factor (TF)–target relationships across species. Despite strong conservation of coding sequences, substantial divergence in regulatory sequences between cultivated and wild rice accessions was associated with distinct regulatory modules and transcriptional programs. Together, these analyses identified gene regulatory networks and transcription factors that contribute to the distinct developmental and photosynthetic features observed in the photosynthetically efficient wild rice species. Gene silencing experiments confirmed the conserved and accession-dependent photosynthetic functions of two candidate regulators, OsDOF8 and OsARID2, identified from gene regulatory networks. This integrative analysis provides a comprehensive transcriptional framework for understanding the coordination between rice leaf development and photosynthesis and identifies potential regulatory targets to optimize leaf traits and photosynthetic efficiency.

## Results

### Variations in leaf developmental and photosynthetic traits across the selected wild and cultivated rice accessions

Based on previous studies showing photosynthetic differences in flag leaves (Giuliani et al., 2013; Mathan et al., 2021a), we selected four representative rice accessions, two Asian cultivated rice *Oryza sativa* ssp. *japonica* cv. Nipponbare (NIP) and *Oryza sativa* ssp. *indica* cv. IR64 (IR64), the African cultivated rice *Oryza glaberrima* (GLA), and the wild rice *Oryza australiensis* (AUS), which showed distinct morphological and anatomical features at different stages of the leaf developmental gradient (Supplementary Figs. 1a-c). Wild rice AUS had a larger Shoot Apical Meristem (SAM) compared with cultivated accessions (Figs. 1a-c). AUS also showed increased cell division activity and an extended division zone, as well as longer epidermal cells at the leaf tip than *O. sativa* accessions across P3 to P5 stages (Figs. 1d, e). Together, increased cell division and elongation contribute to the longer leaf primordia in AUS compared with cultivated accessions (Fig. 1f). AUS also showed significantly larger stomata compared to the cultivated accession at P5, while stomata number remained comparable among the accessions (Figs. 1g, h).

**Figure 1.**
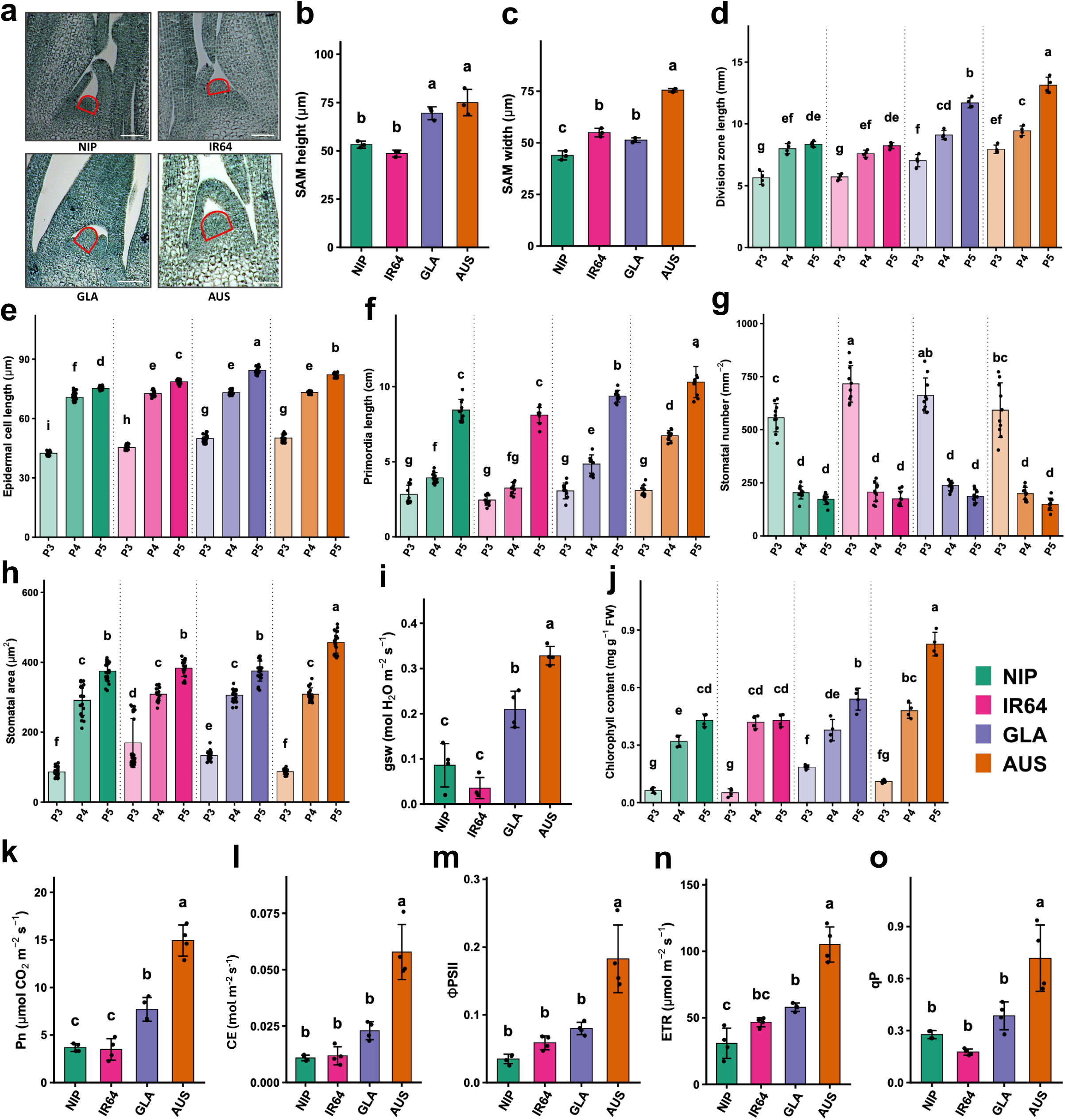
Leaf developmental and photosynthetic differences across selected wild and cultivated rice accessions. **(a)** Representative longitudinal sections of the shoot apical meristem and surrounding leaf primordia (SAM+Pi) at one week post-germination in *Oryza sativa* ssp. *japonica* cv. Nipponbare (NIP), *O. sativa* ssp. *indica* cv. IR64 (IR64), *O. glaberrima* (GLA), and *O. australiensis* (AUS). The SAM region is outlined in red. Scale bar = 250 µm. **(b-c)** Quantification of SAM height **(b)** and width **(c)** across the four accessions. **(d–h)** Quantification of division zone length **(d)**, epidermal cell length **(e)**, leaf primordium length **(f)**, stomatal density **(g)**, and stomatal area **(h)** at P3, P4, and P5 stages across the selected accessions. **(i-j)** Stomatal conductance (gsw) **(i)** at P5 stage and chlorophyll content **(j)** at P3, P4, and P5 stages across the selected accessions. **(k–o)** Quantification of net photosynthetic rate (Pn) **(k)**, carboxylation efficiency (CE) **(l)**, effective quantum yield of PSII (ΦPSII) **(m)**, electron transport rate (ETR) **(n)**, and photochemical quenching (qP) **(o)** at the P5 stage of the selected accessions. The lowercase letters indicate significant differences at *P* < 0.05 based on one-way ANOVA followed by Tukey’s HSD test. Accessions are colour-coded as NIP – green, IR64 – magenta, GLA – purple, and AUS – orange, with stage-specific shading used to distinguish the P3, P4, and P5 stages.

In addition to developmental differences, significant variation in photosynthetic traits was evident among the accessions during early leaf development. AUS exhibited significantly higher stomatal conductance (gsw) and higher total chlorophyll content at P5 than cultivated accessions (Fig. 1i, j). The remarkable increase in chlorophyll content from P3 to P4 and P5 was evident across all four accessions. Quantification of biochemical traits, including net photosynthetic CO₂ assimilation rate (Pn) (Fig. 1k) and carboxylation efficiency (Fig. 2l), together with photochemical traits such as effective quantum yield of PSII (ΦPSII) (Fig. 1m), electron transport rate (ETR) (Fig. 1n), and photochemical quenching (qP) (Fig. 1o), showed the higher photosynthetic efficiency of AUS compared with cultivated accessions. Taken together, distinct stage- and species-specific developmental and photosynthetic differences were evident between wild rice AUS and cultivated accessions along the leaf developmental gradient.

**Figure 2.**
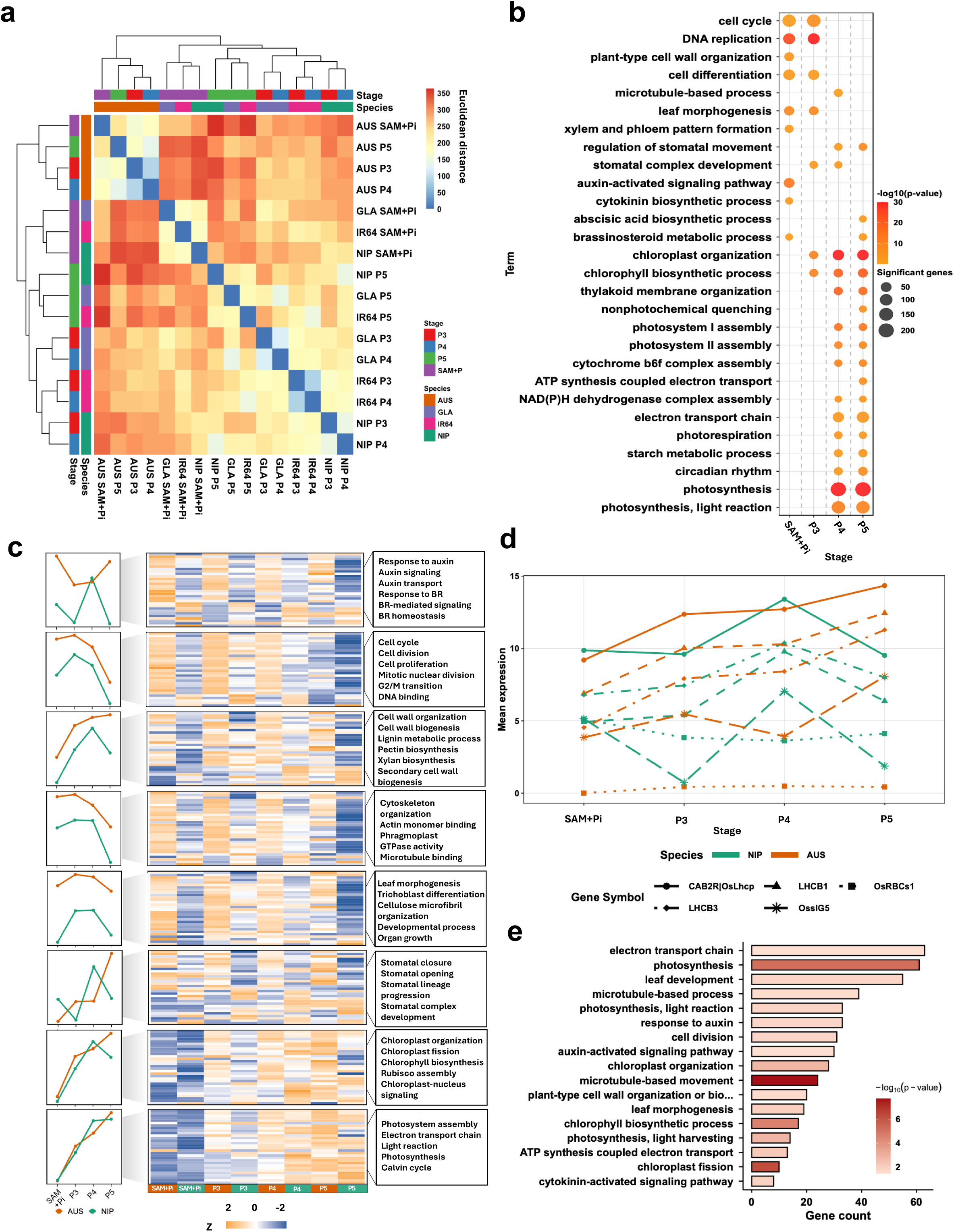
Variations in gene expression and associated biological processes across different leaf developmental stages of the selected rice accessions. **(a)** Heatmap showing Euclidean distances for gene expression profiles, calculated from log₂-transformed normalized counts and grouped by accession and developmental stage. Annotation bars indicate accession and stage. **(b)** Bubble plot showing enriched Gene Ontology (GO) terms for genes predominantly expressed in SAM+Pi, P3, P4, and P5 developmental stages. Bubble size represents the number of significant genes associated with each term, and color indicates enrichment significance as −log₁₀(*P*-value). **(c)** Dynamic expression pattern of prominent pathway-specific differentially expressed gene (DEG) sets associated with developmental and photosynthetic processes across different stages in *O. australiensis* (AUS) and *Oryza sativa* ssp. *japonica* cv. Nipponbare (NIP). The line plots show mean expression trajectories for each DEG set, while the heatmap shows row-wise Z-scores calculated from log₂-transformed normalized expression values. Associated enriched GO terms are shown in the right panel. **(d)** Expression profile of the top five photosynthesis-related DEGs across developmental stages in AUS and NIP. Values represent mean log₂-transformed normalized expression, with individual genes shown as lines and coloured by accession. **(e)** GO terms enriched among genes dynamically upregulated across developmental stages in AUS relative to NIP. Bar length represents gene count, and colour intensity indicates enrichment significance as −log₁₀(*P*-value).

### Gene expression profile and associated biological processes along the leaf developmental gradient of the selected rice accessions

Given the striking developmental and photosynthetic differences across leaf primordia stages, we investigated the transcriptional landscape along the leaf developmental gradient for the selected accessions. Gene expression profiles across developmental stages were distinct in wild rice AUS compared with the cultivated rice accessions (Supplementary Figs. 2a-e, Supplementary Table 1). Transcriptional changes along the leaf developmental gradient revealed more differentially expressed genes (DEGs) between early proliferative SAM+Pi and late photosynthetic P5 stages compared to the P3 and P4 stages for each species (Supplementary Figs. 2f-i, Supplementary Table 2, Supplementary Dataset S1). We then quantified the relative contributions of developmental stages and species to global transcriptomic variation using non-parametric PERMANOVA (Permutational Multivariate Analysis of Variance) on the Euclidean distance matrix (Anderson, 2014). The combined model (Stage + Species) explained 82.5% of the total variance (p < 0.001), with species accounting for a larger proportion (R² = 50.7%, F = 39.68, p < 0.001) than stage (R² = 31.8%, F = 24.89, p < 0.001). The largest species-level separation was observed between AUS and NIP (Supplementary Fig. 2j), supporting distinct transcriptional landscapes between the wild and cultivated rice. Among the leaf developmental stages, SAM+Pi and P5 were the most divergent, while P3 and P4 were the most similar (Supplementary Fig. 2k, Supplementary Dataset S1). Analysis of Species × Stage centroid distances further distinguished cross-stage interspecific differences. AUS at SAM+Pi and NIP at P5 were transcriptionally most divergent, whereas IR64 P3 and P4 were the most similar, reflecting the combined effects of developmental progression and species divergence (Fig. 2a). Stage-specific transcriptional signatures showed the highest conservation at SAM+Pi, followed by P5. The P3 and P4 stages showed relatively fewer shared genes among accessions, indicating species-specific transcriptional profiles for transitional stages (Supplementary Fig. 3; Supplementary Datasets S2 & S3).

Detailed analyses of DEGs and stage-specific signatures identified key genes and biological processes underlying the leaf developmental and photosynthetic progression (Supplementary Figs. 4 & 5; Supplementary Datasets S2 & S3). Genes involved in SAM maintenance and activity, leaf patterning and vascular development were preferentially expressed in SAM+Pi. In contrast, genes associated with auxin signaling, cell cycle progression, and morphogenesis showed higher expression in both SAM+Pi and P3 across accessions (Supplementary Figs. 4 & 5). Consistently, Gene Ontology (GO) analysis of genes upregulated at these stages showed predominant enrichment for developmental processes (Fig. 2b; Supplementary Figs. 6a, b; Supplementary Dataset S4). Genes associated with light harvesting, photosystems, and chloroplast organization were expressed at higher levels in the P4 stage, while genes related to the core photosynthetic machinery, including the electron transport chain, Calvin cycle, photorespiration, and Rubisco-associated functions, predominated at P5 across all four accessions (Supplementary Figs. 4c, d & 5). Accordingly, genes highly expressed at P4 and P5 were enriched for photosynthesis-related processes (Fig. 2b; Supplementary Figs. 6c, d; Supplementary Dataset S4). Notably, chlorophyll- and chloroplast-associated terms, along with stomata-development-related processes, began to appear from P3 onward, indicating that the P3–P4 represents a key transition in the establishment of photosynthetic capacity (Fig. 2b; Supplementary Figs. 4b-c; 5 & 6b-c).

Since variation in developmental and photosynthetic traits, together with Euclidean distance analysis, identified AUS and NIP as the most distinct accessions, we examined the dynamic expression profiles of prominent pathway-specific genes across developmental stages in the two accessions. AUS not only showed higher expression of key developmental and photosynthesis-related genes, but also maintained their expression into later developmental stages (Fig. 2c; Supplementary Figs. 7), indicating a sustained developmental-to-photosynthetic transcriptional programme relative to NIP. Consistent with this pattern, expression of the top five photosynthesis-related DEGs continued to increase at P5 in AUS contrary to NIP (Fig. 2d). Similar trends was observed for genes associated stomatal development, chlorophyll biosynthesis, chloroplast organization, and photosystem assembly (Fig. 2c; Supplementary Figs. 7). Genes associated with the cell cycle, cell wall organization, and morphogenesis showed higher expression in AUS than in NIP at all developmental stages. GO enrichment analysis further showed that major developmental and photosynthetic functions, including cell division, leaf development, chlorophyll biosynthesis, chloroplast organization, electron transport chain, and photosynthesis, were enriched for genes dynamically upregulated in AUS compared to NIP across developmental stages (Fig. 2e; Supplementary Fig. 6e). Together, these results indicate that AUS maintains stronger and sustained developmental and photosynthetic transcriptional signatures across the leaf developmental gradient.

### Stage-resolved gene co-expression analysis revealed divergent transcriptional regulation of leaf developmental and photosynthetic progression across different rice accessions

Principal Component Analysis (PCA) combined with Self-organising map (SOM) clustering resolved nine gene expression clusters in each accession that represented either stage-specific or shared expression patterns across leaf developmental stages (Supplementary Figs. 8a–d; Supplementary Datasets S5 & S6). The genes and enriched biological processes within these clusters were consistent with those identified from differential expression analysis, supporting the robustness of the stage-resolved transcriptome patterns (Supplementary Figs. 8e–h; Supplementary Datasets S5 & S6). Weighted Gene Co-expression Network Analysis (WGCNA) using the PCA-SOM-derived gene clusters identified co-expression modules linked to distinct stage- and species-specific biological processes, including early development and morphogenesis in SAM+Pi/P3-associated modules and photosynthetic establishment and function in later P4/P5-associated modules (Fig. 3; Supplementary Dataset S7). Despite broader conservation of stage-associated biological processes, limited overlap was observed for module gene composition among accessions, with AUS particularly showing distinct stage-resolved expression profiles (Supplementary Fig. 9a). Hub transcription factors (TFs) composition was similarly divergent, with limited overlap for the top 25 hub TFs in stage-specific modules among accessions (Supplementary Figs. 9b-i & 10). This divergence was especially evident in modules associated with morphogenesis and photosynthetic establishment. TF families, such as MYB/SANT, Homeodomain, CO-like, NAC/NAM, etc., appeared as hub TFs in AUS. In contrast, NIP showed preferential representation of bHLH, MADS box, Sigma 70, GATA, etc. hub TF families as key regulators (Fig. 3e; Supplementary Dataset S8). Together, these analyses indicate that although the biological processes underlying leaf development and photosynthetic progression are broadly conserved, the co-expression architecture and putative regulatory TFs are substantially rewired.

**Figure 3.**
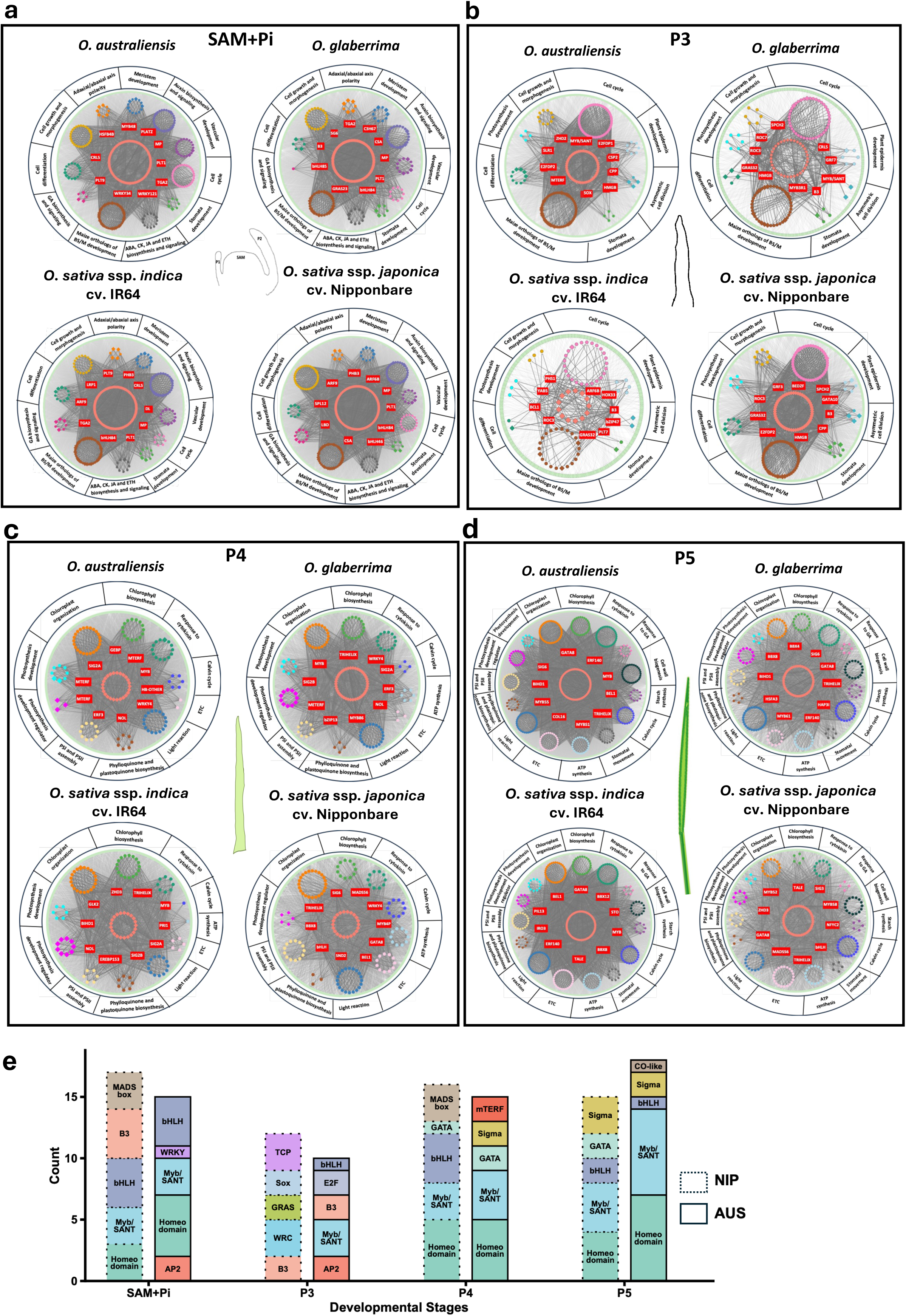
Stage-specific co-expression networks identify candidate regulators of leaf development and photosynthesis. **(a–d)** WGCNA-derived gene co-expression networks for SAM+Pi **(a)**, P3 **(b)**, P4 **(c)**, and P5 **(d)** stages across *O. australiensis*, *O. glaberrima*, *O. sativa* ssp. *indica* cv. IR64, and *Oryza sativa* ssp. *japonica* cv. Nipponbare. The networks show stage-specific co-expression modules associated with leaf development and photosynthesis. The central coral circle indicates the hub transcription factor (TF) node, and the top 10 hub TFs are highlighted in red boxes. Coloured subnetworks represent genes grouped by annotated biological processes, shown in the outer ring. Within each subnetwork, nodes represent genes or TFs associated with that process, whereas genes not assigned to a major annotated process are placed in the surrounding pale green region. Dark edges indicate connections between hub TFs and genes within annotated subnetworks; lighter edges represent connections to other co-expressed genes outside these groups. **(e)** Stacked bar plots showing distribution of TF families among the top 25 hub TFs in *O. australiensis* (AUS) and *Oryza sativa* ssp. *japonica* cv. Nipponbare (NIP) across developmental stages. Bar height represents the number of TFs in each family, and colors denote different TF families.

### Synteny-anchored orthology analysis revealed promoter-level regulatory divergence between cultivated and wild rice

We next examined whether variations in module composition and transcriptional regulation across the accessions along the leaf developmental gradient were associated with regulatory sequence divergence among orthologous genes. All-vs-all synteny analysis for orthologous genes, inferred from protein-level orthology, showed that 76.72% of the analyzed genes were involved in at least one collinear relationship, indicating extensive conservation of coding-gene collinearity (Supplementary Dataset S9). Genes involved in chlorophyll biosynthesis, chloroplast organization, light-harvesting, the Calvin cycle, and other photosynthetic processes were generally retained within conserved syntenic contexts across the four rice accessions. Noteworthy was the conserved syntenic context of the ten most dynamic photosynthesis-related DEGs, including *OsCAB1R*, *OsGAPDH*, *OsHPR1*, *OsPsBW*, *OsHOX16,* and *OsLHCB4* (Fig. 4a). Despite this strong collinearity, their regulatory sequences were substantially divergent, particularly for the wild rice AUS. Similarly, dynamic developmental genes associated with phytohormone signalling, cell-wall modification, cell-cycle progression, morphogenesis, and stomatal development were also represented within collinear regions, but showed stronger promoter sequence variation in AUS (Supplementary Fig. 11).

**Figure 4.**
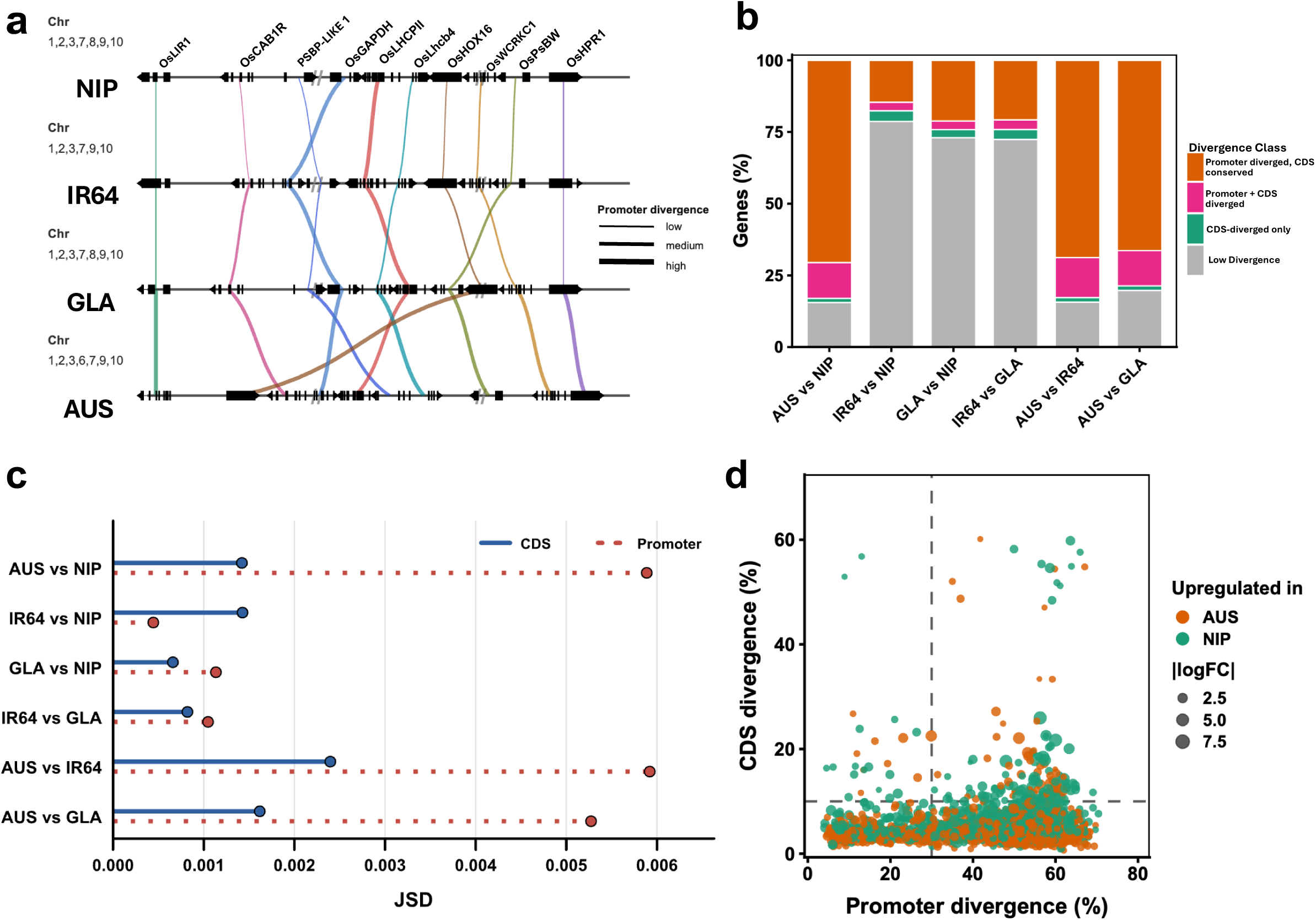
Conserved microsynteny and diverged regulatory sequences for dynamic DEGs across the four *Oryza* genomes. **(a)** Microsyntenic organization of the top ten dynamic photosynthesis-related DEGs across *Oryza sativa* ssp. *japonica* cv. Nipponbare (NIP), *O. sativa* ssp. *indica* cv. IR64 (IR64), *O. glaberrima* (GLA), and *O. australiensis* (AUS). Gene names are shown above the NIP reference track, with chromosome numbers indicated on the left. Horizontal tracks represent genomic intervals, and gene models are displayed below the corresponding labels, with individual blocks representing exons. Coloured links connect collinear gene pairs across genomes, with line thickness representing divergence within the 2-kb upstream regulatory region; thicker lines indicate greater divergence at the promoter. **(b)** Pairwise distribution pattern for divergence of coding DNA sequence (CDS) and promoter sequence across the four genomes. Genes were assigned to four classes: low divergence, CDS-specific divergence, combined promoter and CDS divergence, and promoter-diverged with CDS-conserved. Stacked bars show the proportional representation of each class across pairwise genome comparisons. **(c)** Pairwise comparison of k-mer divergence in CDS and promoter sequences among the four rice accessions using Jensen–Shannon divergence (JSD). Solid blue lines indicate CDS divergence, whereas dotted red lines indicate promoter divergence on the same JSD scale. **(d)** Representation of CDS and promoter sequence divergence for developmental and photosynthetic DEGs between NIP and AUS. Each dot represents a DEG, coloured by the accession in which the gene is upregulated. The dot size indicates the absolute log₂ fold change. Dashed lines indicate the thresholds to define low- and high-divergence classes.

Promoter sequences of orthologous genes showed substantial divergence between AUS and NIP, with an average percent identity (PID) of 53.22% (Supplementary Dataset S10). The promoter divergence was observed despite 92.56% conservation of coding DNA sequences (CDS) between NIP and AUS (Supplementary Dataset S10). Accordingly, the proportion of genes with higher promoter divergence was substantially greater in comparisons involving AUS than in comparisons among the cultivated accessions (Fig. 4b; Supplementary Figs. 12a, b). Consistent with this global promoter variation pattern, cross-accession k-mer distribution analysis showed greater regulatory-sequence divergence involving AUS compared with the cultivated accessions (Fig. 4c). Jensen–Shannon divergence showed a strong promoter sequence divergence between AUS and NIP (JSD = 5.89 × 10⁻³), indicating structurally distinct AUS promoters compared with cultivated accessions. Moreover, dynamic expression of development- and photosynthesis-related DEGs showed a stronger association with promoter divergence than with CDS divergence (Fig. 4d; Supplementary Figs. 12c, d). Together, these results suggest that conserved coding loci involved in leaf development and photosynthesis harbour divergent upstream regulatory sequences, particularly in AUS, which may contribute to accession-specific variations in expression profiles across the leaf developmental gradient.

### Transcriptional regulators of development and photosynthetic efficiency along the rice leaf developmental gradient

Given the observed promoter sequence variation among the four accessions, we next examined whether stage-specific gene co-expression modules were associated with divergent regulatory interactions. Gene regulatory networks (GRNs) were independently inferred for each accession using stage-specific WGCNA modules and the corresponding accession-specific promoter sequences. TF–target relationships were predicted using GENIE3, a random forest-based approach, where candidate regulators for each target gene were restricted to transcription factors with cognate binding motifs detected in the corresponding promoter region (FDR < 0.1; Supplementary Dataset S11; Huynh-Thu et al., 2010). TF–target regulatory networks were then integrated across species and developmental stages and functionally annotated for the developmental, transitional, and photosynthetic processes and pathways (Tian et al., 2020) (Supplementary Datasets S12 & S13). Based on network topology, stage and accession coverage, and the distribution of targets for desired biological pathways across species and developmental stages, TFs were classified into “Core”, “Multifunctional”, and “Category-specific” TFs. “Core” TFs formed a densely connected and broadly conserved network layer, targeting genes across multiple developmental and photosynthetic pathways (Fig. 5a). “Multifunctional” TFs displayed intermediate to high connectivity, linked distinct regulatory modules, and targeted both developmental and photosynthetic genes, indicating roles in coordinating leaf development with the establishment photosynthetic capacity (Fig. 5b). In contrast, “Category-specific” TFs showed functional specificity of target genes for either development or photosynthetic transition and efficiency (Supplementary Fig. 13). AP2 and DOF family members were prominent among the “Core” regulators, whose predicted targets spanned cell-cycle regulation, cell wall-associated functions, and auxin signaling, as well as chlorophyll biosynthesis, chloroplast organization, the Calvin cycle, light reactions, and electron transport (Fig. 5a; Supplementary Figs. 14a, b). Consistent with the broader regulatory association, approximately 45% of the annotated core DOF targets were development-related, while approximately 41% were associated with photosynthesis (Supplementary Datasets S11 & S12). Members of MYB/SANT, AP2, NAC/NAM, TCR/CxC, and TCP TF families were largely represented in “Multifunctional” TFs that were predicted to regulate genes involved in early developmental programs to the progressive establishment and maintenance of photosynthetic capacity (Fig. 5b, c; Supplementary Figs. 14). Among the highly variable “Category-specific” TFs, members of the MYB/SANT, TCP, and Homeodomain TF families were identified as major regulators of developmental genes. In contrast, bZIP, bHLH, WRKY, AP2 and NAC/NAM TF families were found to target genes involved in cellular differentiation and photosynthetic transition in leaves (Fig. 5d; Supplementary Figs. 13 & 14). Members of the bZIP, B3, BES1 C2H2 ZF, HSF, and MADS box TF families were identified as important regulators of photosynthetic genes (Fig. 5d; Supplementary Fig. 13). Together, these results indicate a layered regulatory architecture comprising conserved core regulators, multifunctional TFs that control developmental and photosynthetic progression, and category-specific TFs with more restricted biological functions.

**Figure 5.**
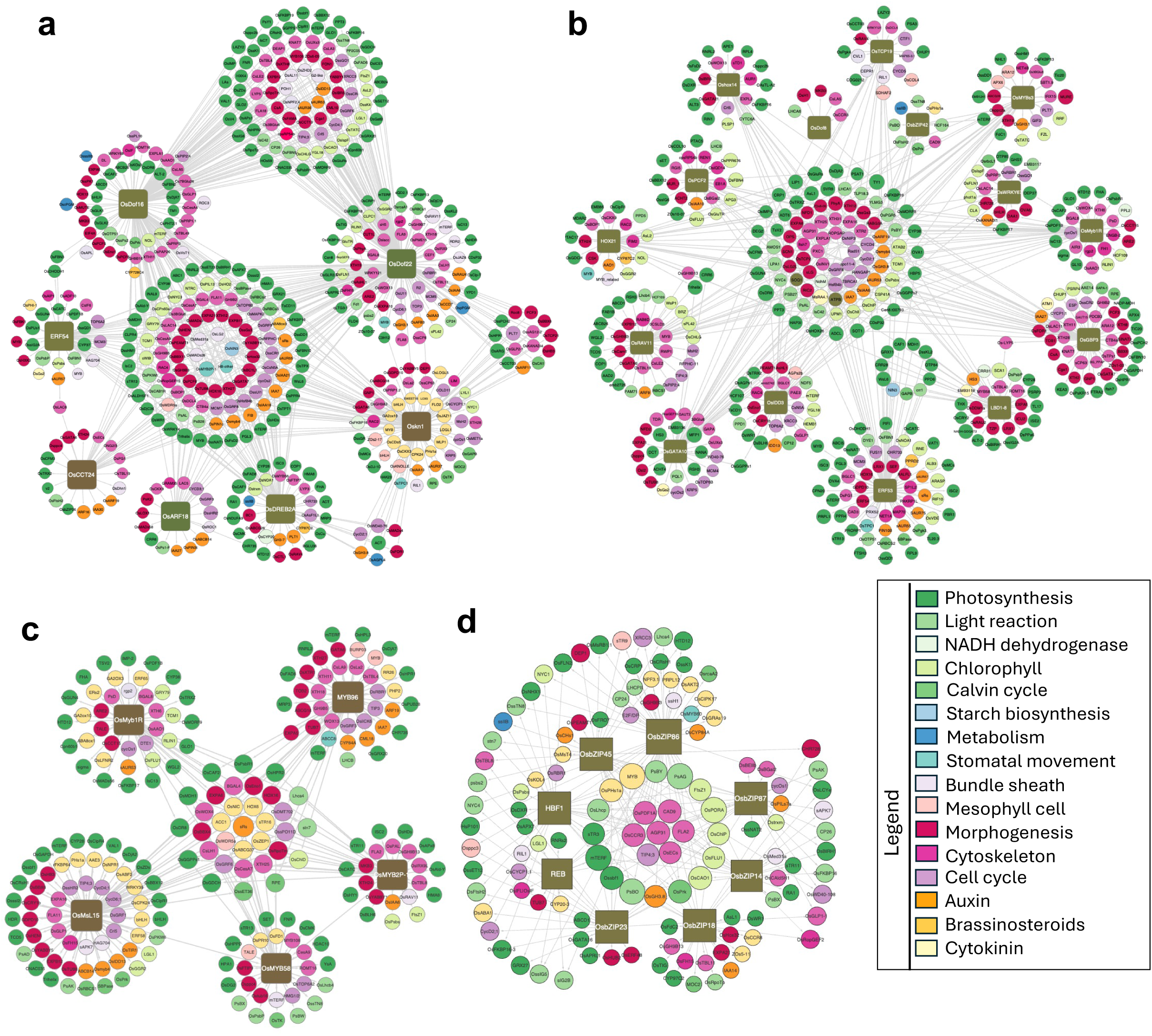
Gene regulatory networks of selected “core” and “multifunctional” transcription factors conserved across the four rice accessions. **(a)** Gene regulatory network of candidate “core” transcription factors (TFs) and their conserved predicted targets across *O. australiensis*, *O. glaberrima*, *O. sativa* ssp. *indica* cv. IR64, and *Oryza sativa* ssp. *japonica* cv. Nipponbare. **(b)** Gene regulatory network of candidate “multifunctional” TFs and their conserved predicted target genes across the majority of accessions, highlighting TFs connected to multiple functional categories. **(c–d)** Gene regulatory subnetworks showing conserved predicted targets of MYB/SANT TF family members **(c)** and bZIP TF family members **(d)** across the four accessions. Only regulatory edges conserved across all four accessions are shown. Node colors indicate the functional categories of target genes, while node size denotes number of co-regulating TFs.

### Rewiring of regulatory networks of key transcription factors and their targets across the four accessions

To examine the contribution of regulatory sequence variations to differential TF–target connectivity and regulatory rewiring across TF classes and accessions, we performed a detailed comparison of TF–target regulatory networks across the four accessions. Only a limited proportion of regulatory edges was conserved for “Core” TFs (∼24.2%) and “Multifunctional” TFs (∼21.33%) across all accessions (Supplementary Dataset S14). Regulatory edge conservation was similarly low for “Category-specific” TFs associated with development (∼18.13%), photosynthetic transition (∼17.61%), and photosynthetic efficiency (∼21.8%) (Supplementary Dataset S14), indicating extensive regulatory rewiring across accessions. Consistent with this, the number of unique targets, along with the associated biological processes regulated by individual TFs, also varied across species (Supplementary Dataset S14). For example, OsARF11 was predicted to target more genes associated with the cell cycle, cell wall, and photosynthesis in AUS than NIP (Fig. 6a), while OsPIL13 showed a greater number of targets for chlorophyll- and chloroplast-related genes in AUS compared to cultivated varieties (Fig. 6b). Some of TFs, such as OsWRKY71, showed functional rewiring across accessions with a comparable number of targets associated with different biological processes (Supplementary Fig. 15a). Temporal rewiring was also evident, for example *OsERF47* activity was predominant at the SAM+Pi in AUS but shifted towards P5 in cultivated *O. sativa* accessions (Supplementary Fig. 15b). Together, these illustrate stage- and accession-dependent rewiring of regulatory networks of key TFs (Fig. 6a-b; Supplementary Fig. 15).

**Figure 6.**
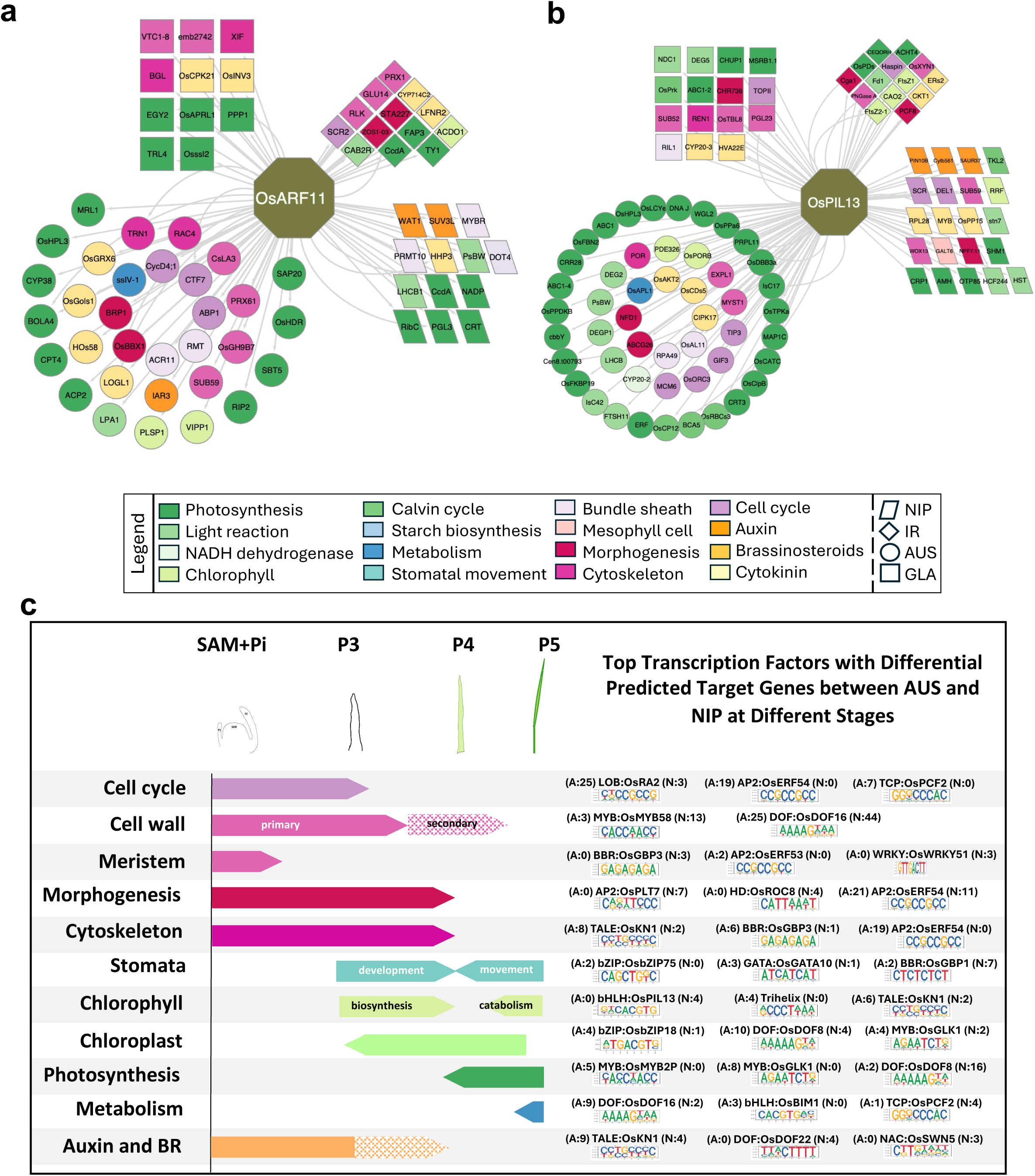
Species-specific regulatory rewiring of key transcription factors across the selected rice accessions. **(a–b)** Gene regulatory subnetworks showing predicted accession-specific target genes of *OsARF11* **(a)** and *OsPIL13* **(b)** across *Oryza sativa* ssp. *japonica* cv. Nipponbare (NIP), *O. sativa* ssp. *indica* cv. IR64 (IR64), *O. glaberrima* (GLA), and *O. australiensis* (AUS). Node colors indicate functional pathway categories of target genes, and node shapes denote accessions. **(c)** Differential regulation of key biological processes at different leaf developmental stages by TFs targeting different target genes between AUS and NIP. Stage-specific pathways are shown alongside TFs predicted to differentially regulate pathway-associated genes. Numbers indicate the number of genes targeted by the TF in the respective accession: A – AUS; N – NIP.

We next examined stage-specific gene regulatory signatures to dissect how key TFs regulate desired biological pathways across accessions. Distinct differences in regulatory patterns of TF families were observed between wild rice AUS and NIP. TF families such as MYB/SANT, B3, TCP, DOF, and NAC/NAM were predicted to regulate a substantially higher number of cell cycle–related genes in SAM+Pi of AUS (Fig. 6c). For instance, *OsERF54* and *OsPCF2* were associated with at least 19 and 7 unique regulatory interactions, respectively, with cell-cycle related genes in AUS compared with NIP (Fig. 6c). Similarly, genes related to meristem development, cell wall organization, morphogenesis, and auxin and brassinosteroid pathways were differentially regulated at SAM+P*_i_* and P3 stages between AUS and NIP by DOF, MYB/SANT, BBR, LOB, GATA, B3, and TCP TF families (Fig. 6c). Moreover, the comparison also identified TFs, such as OsERF53/54, OsDOF16/18, OsGATA6/10, OsbZIP75, and OsGBP1, that showed differential regulation of genes involved stomatal development between the wild and the cultivated rice (Fig. 6c). TF families, including Trihelix, bZIP, bHLH, AP2, MYB/SANT, TALE, and WRKY showed notable differences in regulation of genes involved in chlorophyll biosynthesis and chloroplast organization (Fig. 6c). Furthermore, NAC, AP2, DOF, GATA, BES, and HSF TF families were found to differentially regulate photosynthesis-associated genes between AUS and NIP (Fig. 6c). Interestingly, the analysis also detected the TFs, such as OsbZIP18 and OsGLK1, with a markedly higher number of uniquely regulated photosynthesis-related genes in wild rice AUS (Fig. 6c) and OsDOF8 with a greater number of photosynthesis-related targets in the cultivated rice NIP (Fig. 7a).

**Figure 7.**
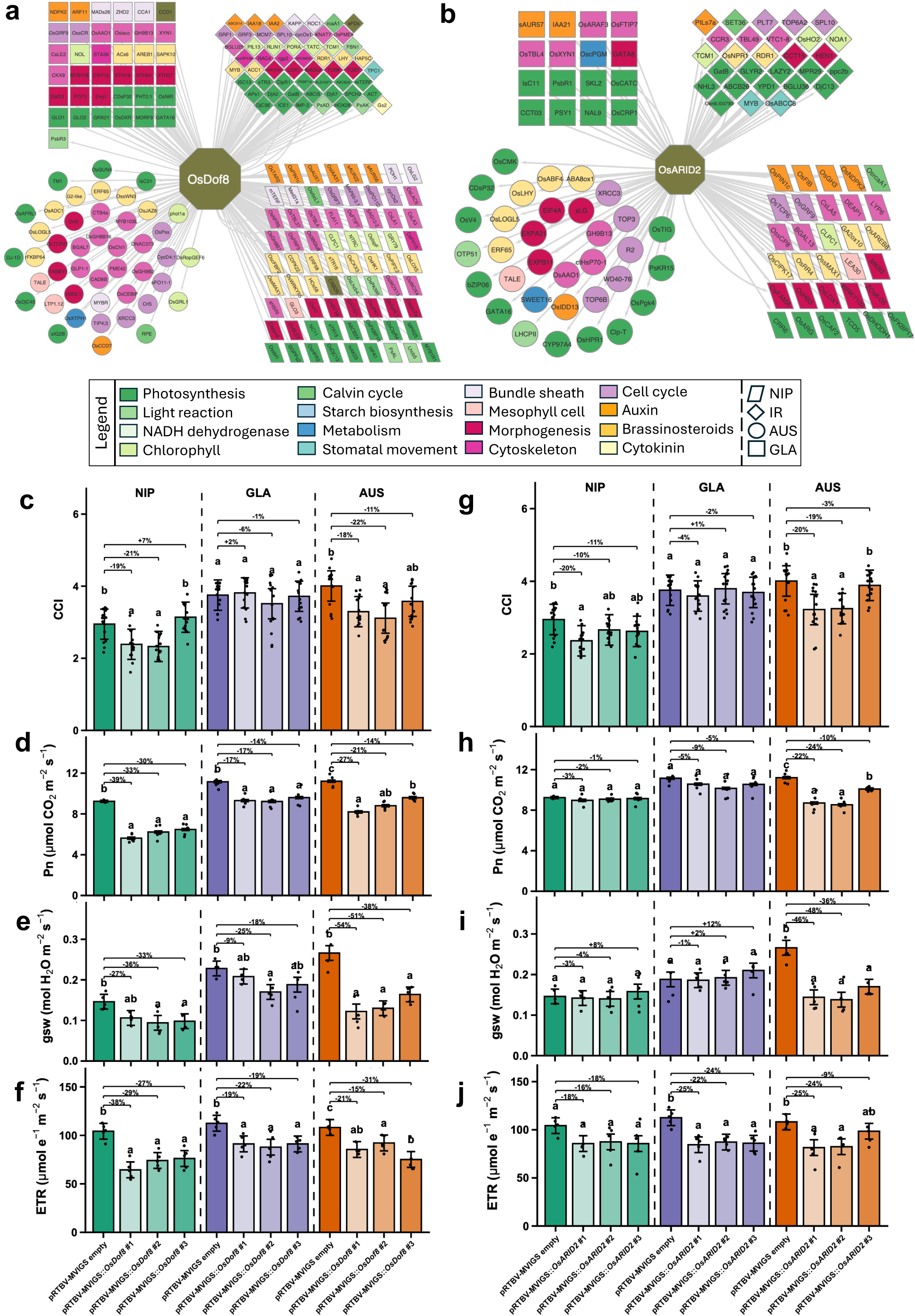
OsDOF8 and OsARID2 links predicted regulatory subnetworks with photosynthetic trait variation across rice accessions. **(a-b)** Gene regulatory subnetworks showing predicted accession-specific target genes for *OsDOF8* (a) and *OsARID2* (b) with candidate target genes across functional pathway categories. Node colors indicate functional pathway categories, and node shapes denote accession. **(c–f)** Quantification of chlorophyll content index (CCI) **(c)**, net photosynthetic rate (Pn) **(d)**, stomatal conductance (gsw) **(e)**, and electron transport rate (ETR) **(f)** in *OsDOF8*-silenced *Oryza sativa* ssp. *japonica* cv. Nipponbare (NIP), *O. glaberrima* (GLA), and *O. australiensis* (AUS) plants. **(g–j)** Quantification of chlorophyll content index (CCI) **(g)**, net photosynthetic rate (Pn) **(h)**, stomatal conductance (gsw) **(i)**, and electron transport rate (ETR) **(j)** in *OsARID2*-silenced NIP, GLA, and AUS plants. Statistical comparisons were performed using one-way ANOVA followed by Tukey’s HSD test; different letters indicate significant differences at *P* < 0.05. Empty-vector plants were used as controls. Percent values above bars indicate reduction relative to the corresponding empty-vector control. Accessions are color-coded: NIP – green, GLA – purple, and AUS – orange, with shading indicating silencing events.

### Silencing the TFs identified from regulatory networks showed accession-specific effects on photosynthetic efficiency

Based on their predicted regulatory properties, two TFs were selected to assess their accession-dependent contribution to photosynthetic regulation. OsDOF8 emerged as a broadly conserved “Multifunctional” regulator with photosynthesis-related genes accounting for 51% of its annotated candidate target genes (Fig. 7a). In contrast, OsARID2 was predicted to act in a more accession-specific manner, showing more than 92.7% cross-accession variation in its predicted target genes (Fig. 7b). The predicted targets of OsDOF8 included genes involved in chlorophyll biosynthesis, chloroplast organization, and light harvesting and photosystem assembly. *OsDOF8* was also predicted to regulate genes associated with cell cycle progression, cell wall formation, morphogenesis, and hormone signaling. Notably, *OsDOF8* showed unique regulatory associations with several genes linked to mature chloroplast function, photosystem organization, light signaling, and photosynthetic optimization in NIP (Fig. 7a). These associations suggested that *OsDOF8* may play a stronger role in coordinating photosynthetic performance in NIP. In contrast, *OsARID2* showed a stronger AUS-biased predicted regulatory network and uniquely connected with candidate targets associated with chlorophyll biosynthesis, chloroplast development, photosystem assembly, and light signaling in AUS (Fig. 7b). Based on these predictions, we silenced *OsDOF8* and *OsARID2* across accessions to assess their contribution to photosynthetic efficiency (Supplementary Figs. 16a-b).

Silencing of both TFs altered chlorophyll content and photosynthetic performance, but the magnitude of these effects varied among accessions. *OsDOF8*-silenced NIP plants showed a clear reduction in photosynthetic attributes, including net photosynthetic rate (Pn), ΦCO₂, chlorophyll content, as well as effective quantum yield of PSII (ΦPSII), electron transport rate (ETR), and stomatal conductance (gsw), indicating that OsDOF8 influences both photochemical and gas-exchange components of photosynthesis (Figs. 7c-f; Supplementary Figs. 16c-d). *OsDOF8*-silenced AUS plants showed similar reductions; however, the effects were less pronounced than in NIP (Figs. 7c-f; Supplementary Figs. 16c-d). In contrast, *OsARID2* silencing produced a stronger photosynthetic phenotype in AUS, with reduced chlorophyll content and lower photosynthetic parameters. The corresponding reductions in NIP and GLA were not evident (Figs. 7g-j; Supplementary Figs. 16e-f). Together, these accession-dependent silencing responses supported the predicted regulatory divergence of the two TFs: OsDOF8 appears to act as a more conserved photosynthesis-associated regulator with stronger functional roles in NIP, whereas OsARID2 shows a more AUS-biased role in regulating photosynthetic efficiency.

### Integrative visualization of gene expression and regulatory network landscape along the rice leaf developmental gradient

To facilitate accessibility and exploration of the transcriptome atlas generated in this study, we developed an interactive database, **Rice DEV-LEAF** (**Rice DEV**elopmental Gradient **L**eaf Gene **E**xpression **A**tlas **F**ramework; https://nipgr.ac.in/DEV-LEAF/). Rice DEV-LEAF is a web-based platform designed to enable intuitive exploration of gene expression dynamics during rice leaf development (Supplementary Fig. 17). The database provides normalized gene expression values, orthologous sequence similarity, differential gene expression results, co-expression relationships, genome-wide syntenic blocks, and gene regulatory network information, enabling comparative analysis across the selected accessions and developmental stages. The database was implemented using a Linux–Apache–MySQL– PHP (LAMP) architecture, with datasets and gene annotations stored in a MySQL relational database. The web interface was developed using HTML, CSS, JavaScript, and PHP, allowing dynamic retrieval, visualization, and comparison of gene expression data. PHP scripts process user queries, retrieve relevant data from MySQL tables, and dynamically generate result pages. The database modules allow users to retrieve gene annotations and expression profiles (Locus Search), assess conserved collinear genes and syntenic relationships across accessions (Collinearity Search), examine differential gene expression and associated GO term enrichment (DGE Search), identify genes with correlated expression patterns across developmental stages (Co expression Search), explore transcription factor–target relationships within gene regulatory networks (GRN Search), visualize expression patterns for multiple genes simultaneously (Multi-Gene Search), perform sequence similarity searches against the DEV-LEAF dataset (BLAST Search), and identify constitutively expressed or stage-specific genes (Housekeeping/Tissue-Specific Gene Search). Together, these modules provide an integrated platform for accessing, querying, and analyzing gene expression dynamics along the rice leaf developmental gradient.

## Discussion

Investigating the genetic regulation of leaf developmental and physiological progression and integrating this progression with photosynthetic efficiency is important for optimizing crop performance. The photosynthetic performance is modified through transcriptional regulatory interventions for chloroplast biogenesis and carbon fixation (Wu et al., 2019; Perveen et al., 2020; Li et al., 2020; Yeh et al., 2022; Sanyal and Ranjan, 2026), as well as for leaf cellular traits (Wang et al., 2017; Lawrence et al., 2021; Østerlund et al., 2025). However, the transcriptional programs that coordinate leaf development with the acquisition of photosynthetic competence remain poorly resolved. Comparative analysis of rice accessions, contrasting in developmental and photosynthetic traits, could be instrumental for identifying key regulators that link leaf development and photosynthetic functions to optimize plant performance. For example, wild rice *O. australiensis* exhibits distinct leaf developmental and photosynthetic features compared to cultivated rice accessions from early leaf developmental stages (Fig. 1). A substantial variation in leaf architecture and anatomy was previously documented across the *Oryza* genus (Chatterjee et al., 2016). Similarly, previous studies have identified photosynthetically efficient wild rice accessions, including *O. australiensis*, and attributed higher photosynthetic efficiency to anatomical, biochemical, and photochemical traits (Giuliani et al., 2013; Mathan et al., 2021a). While studies, such as the GA–OsGRF7/8 module mediating leaf growth differences, provided genetic insights into leaf growth differences between cultivated and wild rice accessions (Jathar et al., 2022), the regulatory networks underlying broader developmental and photosynthetic variations remain poorly understood. The distinct transcriptional separation between the wild rice *O. australiensis* and the cultivated accessions revealed by non-parametric PERMANOVA on the Euclidean distance matrix, and the larger contribution of species than developmental stage to transcriptomic variation, suggest that contrasting leaf traits are driven by differential species-specific gene regulatory programs (Fig. 2a; Supplementary Figs. 2).

Stage-specific transcriptional signatures suggest a coordinated progression of leaf development and physiology from meristem maintenance and leaf morphogenesis to photosynthetic maturation across the four accessions. The expression patterns broadly resolved the developmental gradient into three phases. Gene expression profile of SAM+Pi largely overlapped with P3 and was dominated by programs associated with meristem maintenance, cell proliferation, polarity establishment, vascular differentiation, and early leaf patterning, suggesting predominance of developmental progression from SAM to P3 (Fig. 2b). Notably, SAM+Pi showed most conserved signatures across accessions, suggesting that the molecular framework controlling early meristem development is largely maintained across diverse *Oryza* backgrounds (Miya et al., 2021; Sun et al., 2024). The P3–P4 interval defined a second transcriptional phase in which the developing leaf shifted from cellular morphogenesis towards functional photosynthetic differentiation that include activation of stomatal and chloroplast-related programs (Fig. 2b). In agreement with this transcriptional signature and a previous study by van Campen et al., (2016), the sharp increase in chlorophyll content between P3 and P4 across the selected accessions further attests the transition towards photosynthetic competence during this interval (Fig. 1j). However, the limited conservation of the stage-specific transcriptional signatures for P3 and P4 across the selected accessions suggested prominent species-specific regulatory programs during this developmental-photosynthetic transition. In the third phase, the transcriptional signature shifted towards the mature photosynthetic functions with higher expression of genes for photosynthesis at P4 and P5 stages of leaf development (Fig. 2b). Together, the transcriptional signatures support a model in which photosynthetic capacity is established gradually through the coordinated regulation of developmental and physiological programs that attests spatial and temporal partitioning of growth-related and photosynthetic processes in grass leaves (Robertson and Wilkins, 2023; Sun et al., 2024).

Despite broadly comparable developmental and physiological progression across the accessions, the transcriptional signatures associated with leaf development, stomatal differentiation, chloroplast biogenesis, and photosynthesis were maintained at higher levels and for longer in *O. australiensis* than in the cultivated accessions (Fig. 2c). The contrast was especially pronounced for photosynthetic genes that showed continued activation in *O. australiensis* at P5 than in Nipponbare (Figs. 2c & d). The enrichment of developmental and photosynthetic processes among the genes dynamically upregulated in *O. australiensis* may contribute to its enhanced leaf growth and photosynthetic capacity (Figs. 2c & e). Consistent with this, the composition of co-expression modules and hub TFs varied considerably among the accessions, with *O. australiensis* showing strong divergence (Fig. 3). While accession-specific transcriptional signatures were evident, the organization of protein-coding genes remained largely conserved across the four genomes (Figs. 4a-c). Genes associated with leaf development and photosynthesis generally retained comparable genomic contexts across accessions (Fig. 4a). However, microsynteny analyses revealed strong divergence in upstream regulatory sequences between wild rice and cultivated accessions. Notably, genes associated with leaf morphogenesis, stomatal differentiation, chloroplast organisation, and photosynthesis showed stronger promoter-level divergence than coding sequence, particularly in *O. australiensis* (Supplementary Fig. 11). This pattern is consistent with a stable gene-rich core within the *Oryza* genus, despite extensive divergence in structural, repetitive, and regulatory regions in wild *Oryza* genomes, including *O. australiensis* (Stein et al., 2018; Long et al., 2024; Fornasiero et al., 2025; Morrison et al., 2025). Because cis-regulatory polymorphisms and transposable-element insertions upstream of genes can modify transcriptional output (Campbell et al., 2020; Castanera et al., 2023), the pronounced promoter divergence in *O. australiensis* provides a plausible regulatory basis for its distinct developmental and photosynthetic transcriptional signature.

While the promoter-informed regulatory networks resolved a hierarchical transcriptional framework of “Core”, “Multifunctional”, and “Category-specific” TFs (Figs. 5; Supplementary Figs. 13), only 24.2% of regulatory interactions were conserved across the accessions, indicating extensive rewiring of downstream targets across developmental stages and biological processes. For example, OsARF11, known to be involved in meristem function and venation (Sims et al., 2021), showed broader developmental connectivity in *O. australiensis* compared with Nipponbare (Fig. 6a). Similarly, OsGLK1, known to be involved in chloroplast and photosynthetic differentiation (Yeh et al., 2022), had substantially more photosynthesis-associated targets in *O. australiensis* than in Nipponbare (Fig. 6c). In addition to the TFs known to be involved in meristem and photosynthetic functions, we also identified TFs with differential targets across the selected accessions. For example, OsERF54 had several unique cell-cycle targets, and OsGATA10 showed stronger connectivity with stomata-development genes in *O. australiensis* than in Nipponbare (Lim et al., 2024). TFs, such as OsDOF8, were predicted to target chloroplast-organisation genes in *O. australiensis* and mature photosynthetic genes in Nipponbare (Fig. 6c), suggesting rewiring of transcriptional programs involves both a differential number of targets and differential functions across the stages. Silencing of TFs identified from gene regulatory networks provided functional support for this accession-specific rewiring of transcriptional networks (Fig. 7). OsDOF8, which has been shown to activate bundle-sheath-associated promoters and regulate photosynthetic cell identity (Swift et al., 2024), was identified as a conserved “Multifunctional” TF to predominantly target photosynthetic genes across the selected accessions. While *OsDOF8* silencing repressed photosynthetic attributes across the selected accessions, the effect was more pronounced in Nipponbare than in *O. australiensis*, suggesting a stronger contribution of OsDOF8 to photosynthetic regulation in Nipponbare. Conversely, silencing of *OsARID2*, which was predicted to act in a more accession-specific manner, produced a stronger photosynthetic phenotype in *O. australiensis* and limited effects in cultivated accessions. ARID regulators have been associated with basal division and elongation networks, as well as phytohormonal pathways in Nipponbare leaves (Xu et al., 2015; Robertson and Wilkins, 2023). Our results show accession-specific deployment of an ARID regulator for chloroplast and photosynthetic functions.

Together, our study highlights the major events from leaf initiation to photosynthetic competency, delineates the gene regulatory networks, and identifies “Core”, “Multifunctional”, and “Category-specific” regulators for leaf developmental and physiological progression along the rice leaf developmental gradient. The study further provides a genetic framework for the distinctive leaf architecture and superior photosynthetic efficiency of the wild rice *O. australiensis* compared with cultivated varieties and posits this wild rice species as a valuable model for optimizing rice leaf developmental attributes and photosynthetic efficiency. The divergence in upstream regulatory sequences, coupled with the limited conservation of regulatory interactions across accessions and accession-specific effects of silencing candidate TFs for photosynthetic functions, affirms that rewiring of transcriptional networks is a crucial driver of developmental and photosynthetic differences between the wild rice *O. australiensis* and cultivated rice accessions. The comprehensive public database “Rice DEV-LEAF Database (https://nipgr.ac.in/DEV-LEAF/)”, developed from the data and findings from this study, would be a valuable resource for exploring gene expression dynamics, transcription factor functions, and gene regulatory networks along the leaf developmental gradient across the selected wild and cultivated rice accessions.

## Methods

### Plant material and growth conditions

The study utilized four rice genotypes: *Oryza sativa* ssp. *japonica* cv. Nipponbare, *Oryza sativa* ssp*. indica* cv. IR64, the African cultivated rice *Oryza glaberrima* (IR102925), and the wild rice species *Oryza australiensis* (IRGC105272). Seeds were germinated on Petri dishes lined with moist germination paper. Post-germination, seedlings were transferred to a hydroponic system with Murashige and Skoog (MS) medium. The nutrient solution, adjusted to a pH of 5.5, was refreshed every 3 days. Seedlings were grown in a greenhouse with a day temperature of 28°C and a night temperature of 23°C, under a 14-hour light and 10-hour dark cycle.

### Leaf primordia sample collections

After defining the leaf developmental window through microtome sectioning, the fourth leaf at various developmental stages was dissected from the leaf three sheath at the same time of day (around 10:00 AM), based on the morphological age as described by van Campen et al. (2016). Shoot apical meristem with initiating primordia (SAM+Pi) samples were collected from the seedling basal region at the shoot-root junction after removing older leaf sheaths. At the time of collection, primordia3 (P3) measured approximately 1–1.5 cm in *O. sativa* and *O. glaberrima* and 1.5–2 cm in *O. australiensis*, and P4 measured 2.5–3 cm in *O. sativa* and *O. glaberrima* and 4.5–5 cm in *O. australiensis*. P5, measuring 6–7 cm in *O. sativa,* 7–8 cm in *O. glaberrima*, and 10–11 cm in *O. australiensis*, were collected on the second day after the fourth primordium emerged from the leaf three sheath.

### Histology

To investigate leaf development and shoot apical meristem (SAM) size, paraffin sectioning was performed as described by Kladnik (2013). Samples were collected from the seedling base, eight days post-germination, and were fixed in FAA (formaldehyde: glacial acetic acid: ethanol, 1:1:18) for 24 hours at 4°C, followed by dehydration in a graded tert-butanol (TBA) series. Samples were then cleared in xylene and embedded in Paraplast Plus wax. 8-µm-thick sections were cut using a microtome and examined with a phase-contrast microscope. SAM size was quantified using ImageJ software.

### Analysis of primordia length, division zone size, cell length, and stomatal patterning

Primordia length was measured using a ruler, with 10 samples per primordium. For division zone size estimation, primordia were fixed in a 3:1 (v/v) solution of absolute ethanol and acetic acid, then rinsed for 30 minutes in a buffer (50 mM NaCl, 5 mM EDTA, 10 mM Tris-HCl, pH 7). Nuclei were stained in the dark for 2 minutes with 1 mg/ml 4,6-diamidino-2-phenylindole (DAPI) in the same buffer. Fluorescent nuclei were visualized using a Nikon 80i epifluorescence microscope, and division zone size was measured following Sprangers et al. (2016). For cell length profiles and stomatal features, primordia were cleared in absolute ethanol for 48 hours (with one change at 24 hours), then transferred to lactic acid at 4°C. Samples were mounted on slides with glycerol and examined with a phase-contrast microscope at 40x magnification. Cell length and stomatal cell counts were quantified using ImageJ software.

### Measurement of total chlorophyll content

Total chlorophyll content was measured following Porra et al. (1989). Leaf primordia were incubated in 5 mL of ice-cold 80% acetone for 48 hours. After centrifugation at 5000× g for 5 minutes, absorbance was recorded at 663 nm (chlorophyll a) and 645 nm (chlorophyll b). Chlorophyll concentration (a + b) was calculated using (Arnon, 1949) and expressed in µg/mg.

### Quantification of photosynthesis rate and related physiological traits

Photosynthesis was quantified at the widest part of the fully expanded leaf blade at the P5 stage using a LI-6800 portable photosynthesis system, with a Multiphase Flash™ Fluorometer serving as the light source. Quantifications were performed between 9:00 AM and 11:00 AM. The system was set to a constant airflow rate of 500 μmol s⁻¹, a CO_2_ concentration of 400 μmol mol⁻¹, a light irradiance of 1500 μmol m⁻² s⁻¹, and a leaf temperature of 28°C. The chamber humidity was maintained at 70–80% to achieve a leaf-to-air vapor pressure deficit. Six primordia from different plants were analyzed to quantify the net photosynthesis rate (A), intercellular CO_2_ concentration (Ci), stomatal conductance to water (gs), and carboxylation efficiency (CE). Photochemical traits were quantified by measuring steady-state fluorescence (Fs) at the ambient CO_2_ level. Light-adapted maximum fluorescence (Fm′) was obtained by exposing the leaf to a high-intensity light pulse (∼8000 μmol m⁻² s⁻¹ for 0.8 seconds). The minimal fluorescence in the light-adapted state (Fo′) was determined by removing the actinic light and applying a 1-second far-red (735 nm) light pulse.

From these fluorescence measurements, the following parameters were calculated:

- Photosystem II Efficiency (φPSII) = (Fm′ − Fs)/Fm′
- Photochemical Quenching Coefficient (qP) = (Fm′ − Fs)/(Fm′ − Fo′)
- Photosynthetic Electron Transport Rate (ETR) = PFDa × φPSII × 0.5

PFDa represents the absorbed photon flux density, and 0.5 is a correction factor to account for energy partitioning (Maxwell and Johnson, 2000).

### RNA-seq library preparation and sequencing

Total RNA was extracted using TRIzol reagent and purified with DNase I treatment using the Plant Total RNA Kit (Sigma-Aldrich). RNA quality and integrity were assessed using the Agilent 2100 Bioanalyzer (Agilent Technologies). RNA-seq libraries were prepared in triplicate following the protocol described by Townsley et al. (2015). 150-bp paired-end reads were generated after sequencing on an Illumina HiSeq platform.

### Quality filtering, mapping of reads, and differential gene expression analysis

RNA-seq reads were filtered using the NGSQC toolkit (Patel and Jain, 2012) and high-quality RNA-seq reads were aligned to the coding DNA sequences (CDS) of *O. sativa* ssp*. japonica* cv. Nipponbare from the Rice Genome Annotation Project Release 7 (RGAP 7) using Bowtie2 with default settings (Langmead and Salzberg, 2012). The Nipponbare reference was chosen to ensure a uniform reference for cross-accession transcriptomic comparisons. The *O. australiensis* genome, enriched with long terminal repeat (LTR) retrotransposons (∼65% in *O. australiensis* vs. ∼10% in *O. sativa*), exhibits a high degree of conservation of coding regions with *Oryza sativa* (Phillips et al., 2022). Although some *O. australiensis-* and *O. glaberrima*-specific genes absent from *O. sativa* may not have been detected, the workflow effectively captured conserved gene differences that drive leaf development and photosynthetic traits. BAM alignment files, created with SAMtools, were used to extract raw gene counts (Li et al., 2009). These counts were normalized using the Trimmed Mean of M-values (TMM) method (Robinson and Oshlack, 2010). Permutational Multivariate Analysis of Variance (PERMANOVA) was performed on the Euclidean distance matrix derived from transcriptomic profiles using the “**adonis2**” function implemented in the “**vegan**” package in R (Anderson, 2014; Oksanen J et al., 2026). Differential gene expression was performed using “**edgeR”** (Robinson et al., 2010) with a generalized linear model (GLM) and a quasi-likelihood F-test. Genes were considered significantly differentially expressed with a false discovery rate (FDR) below 0.05 and a log fold change (logFC) ≥ 1 or ≤ −1.

### Gene Ontology (GO) enrichment analysis

GO enrichment analysis was performed using GO annotation files (GAF) obtained from the Plant Transcriptional Regulatory Map (PlantRegMap) database and analyzed using the “**topGO”** package in R (Tian et al., 2020; Alexa and Rahnenfuhrer, 2025).

### Principal component analysis with self-organizing map (PCA-SOM) clustering

Normalized read counts were used for PCA-SOM clustering, as described by Chitwood et al. (2013). Clustering was performed to account for both developmental stages within accessions and specific developmental stages across accessions. Genes in the top 50% of the coefficient of variation for expression across developmental stages were used for clustering. Scaled expression values were used to create gene clusters specific to developmental stages within and across accessions, following Wehrens and Buydens (2007). Clustering was performed using a 3×3 hexagonal Self-Organizing Map (SOM) with 100 training iterations. SOM clusters were visualized in PCA space, with principal component (PC) values computed from gene expression data using the “**prcomp**” R package. Line graphs and heat maps for each cluster were generated with the “**superheat**” R package.

### Gene co-expression network analysis

Stage-specific cluster transcripts were used to construct an unsigned correlation network for each developmental stage of the rice accessions using the Weighted Gene Co-expression Network Analysis (WGCNA) package (version 1.68; Langfelder and Horvath, 2008). The data were normalized using z-scores with the R scale function. A soft-thresholding power was selected to achieve a scale-free topology with a fit index of 0.7. An adjacency matrix was generated and subsequently converted into a topological overlap matrix (TOM). Network properties, such as node strength, were derived from the TOM, and the network was visualized using Cytoscape (Gustavsen et al., 2019).

### Collinearity analysis and extraction of species-specific regulatory sequences

Reference genome FASTA and GFF3 annotation files of *Oryza sativa* ssp. *japonica* cv. Nipponbare, *O. sativa* ssp. *indica* cv. IR64, and *O. glaberrima* were obtained from EnsemblPlants (Yates et al., 2022), while the *O. australiensis* genome was sourced from Long et al. (2024). Genome-scale promoter datasets were generated using protein-level orthology analyses followed by coordinate-based extraction of upstream sequence (Emms and Kelly, 2019). Protein-coding genes were first filtered to remove short coding sequences (<300 bp) to minimise low-complexity alignments and spurious similarity matches. Synteny and microsynteny analyses were performed using DIAMOND “blastp” protein similarity searches followed by MCScanX to identify collinear blocks and anchor gene pairs across the four rice accessions (Wang et al., 2012; Buchfink et al., 2014). Orthologous gene pairs were inferred using BLAST-based best-hit criteria under defined thresholds for sequence identity, alignment length, and query coverage (Altschul et al., 1997; Moreno-Hagelsieb and Latimer, 2008; Emms and Kelly, 2019). Chromosome-level FASTA files were indexed before sequence retrieval, and strand-specific 2-kb upstream regions relative to the transcription start site were extracted for all annotated genes. Boundary corrections were applied for loci located near scaffold ends to avoid negative coordinates or intervals extending beyond sequence length. To quantify global CDS and promoter compositional similarity, 8-mer frequency profiles were computed for all orthologous promoter sequences and aggregated for each accession. Pairwise Jensen-Shannon divergence (JSD) was then calculated between accessions using the normalized 8-mer frequency distributions (Lin, 1991).

### Gene regulatory network prediction

The gene regulatory network (GRN) was predicted using GENIE3 (Huynh-Thu et al., 2010), incorporating developmental stage-specific transcripts and hub transcription factors (TFs) identified from the gene co-expression network (GCN) for each developmental stage. FIMO was to scan the 2 kb promoter regions upstream of each transcription start site to identify TF binding motifs (TFBMs) (Grant et al., 2011). TFBM data were obtained from CIS-BP, which includes experimentally determined and inferred motifs for *Oryza sativa*. TF annotations were sourced from CIS-BP (Weirauch et al., 2014) and PlantTFDB (Jin et al., 2017). Transcription factors with TFBMs in the promoter regions were used as predictors in GENIE3’s random forest algorithm. The ranked TF-target interactions generated by GENIE3 were then used to construct the regulatory network using igraph (Csardi and Nepusz, 2006) and visualized in Cytoscape (Gustavsen et al., 2019).

### Virus-induced gene silencing (VIGS) of *OsDOF8* and *OsARID2*

VIGS constructs targeting *OsDOF8* and *OsARID2* were generated using the pRTBV-mVIGS vector following the protocol described by Kant and Dasgupta (2017). Recombinant constructs were introduced into *Agrobacterium tumefaciens* strain EHA105. Bacterial cultures were grown overnight at 28 °C and subsequently subcultured to an OD600 of 0.6–0.8 in the presence of 200 µM acetosyringone. Cells were harvested by centrifugation and resuspended in infiltration buffer containing 10 mM MES, 10 mM MgCl₂, and 200 µM acetosyringone. Approximately 100 µl of bacterial suspension was injected into the meristematic region at the shoot–root junction of 2-week-old rice seedlings using a vertically positioned syringe (Kant et al., 2015). Following inoculation, seedlings were placed on sterile Whatman No. 1 filter paper soaked in MS medium. Roots were covered with moist tissue to prevent desiccation, and seedlings were transferred to MS medium 24 h post-inoculation and maintained at 28 °C under previously described growth conditions.

### RNA extraction and gene expression studies

Seedling leaves were excised using microscissors, immediately frozen in liquid nitrogen, and stored at −80 °C. Total RNA was extracted using QIAzol lysis reagent (Qiagen), and 2.0 µg of total RNA was used for cDNA synthesis with the Verso cDNA Synthesis Kit (Thermo Fisher Scientific). For quantitative PCR (qPCR), cDNA was diluted with nuclease-free water and used as the template. Reactions were performed using gene-specific primers (Supplementary Data S14) and PowerUp™ SYBR™ Green Master Mix (Applied Biosystems) on a Bio-Rad CFX9 Connect real-time PCR system (Bio-Rad). Transcript abundance was normalized to the rice reference genes 18S and/or UBQ, and relative expression levels were calculated using the 2^−ΔCT method (Schmittgen and Livak, 2008). Each experiment included three biological replicates, each with ∼5 technical replicates (n∼15).

## Author contributions

**Conceptualization:** V.J., M.K.P., and A.R. **Methodology:** M.K.P., V.J., and A.R. **Investigation:** M.K.P., V.J., R.S., A.D., R.R. and A.V.D. **Visualization:** M.K.P., V.J., A.T.V, A.D., and R.S. **Data curation:** A.T.V, V.J., M.K.P., S.K. and A.R. **Validation:** R.S., M.K.P., and V.J. **Formal analysis:** A.T.V, V.J., and M.K.P. **Writing—original draft:** V.J., M.K.P., and A.R. **Writing—review and editing:** V.J., M.K.P., A.T.V., R.S., A.D.V, A.D., R.R., S.K., and A.R.

## Acknowledgements

V.J. and M.K.P. acknowledge their UGC-SRF and CSIR-SRF fellowships, respectively. We thank NIPGR Central Instrumentation Facility for their support. The seeds of wild rice *Oryza australiensis* and *Oryza glaberrima* were kindly provided by Dr. Kuldeep Singh and Dr. Kumari Neelam, Punjab Agricultural University, Ludhiana, India.

## Availability of data and materials

The data supporting the findings of this study are available in the supplementary materials of this article. The quality-filtered RNAseq reads are deposited to the NCBI Short Read Archive (SRA) under accessions SRR15144739 - SRR15144754. The datasets generated and analysed during this study are also available as a public database “Rice DEV-LEAF (https://nipgr.ac.in/DEV-LEAF/)”.

## Supplementary Figures

**Supplementary Figure 1.**
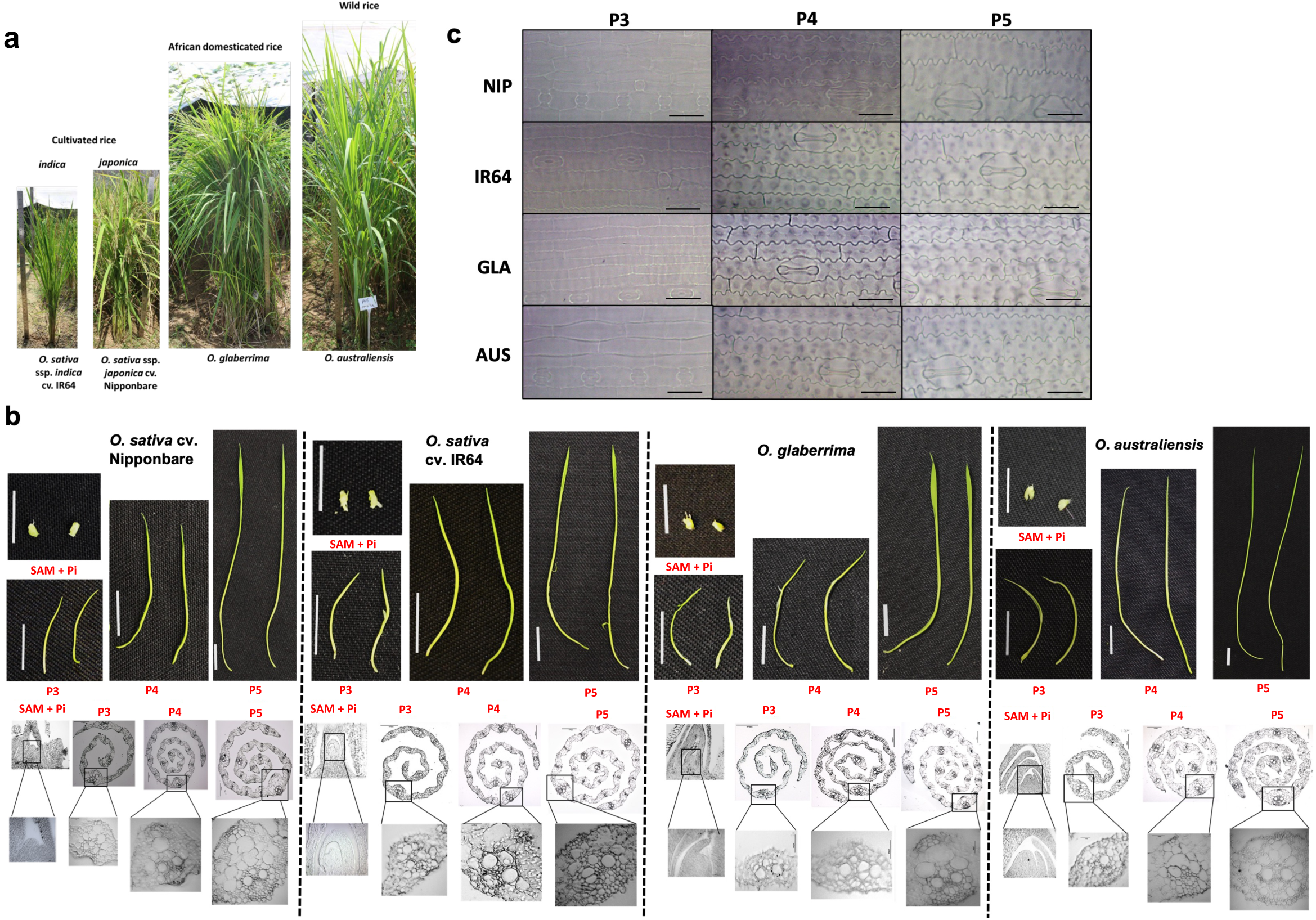
Morphological and anatomical variations across the selected wild and cultivated rice accessions. **(a)** Representative images of the field-grown plants of the four rice accessions used in this study: *Oryza sativa* ssp. *japonica* cv. Nipponbare (NIP), *O. sativa* ssp. *indica* cv. IR64 (IR64), *O. glaberrima*(GLA), and *O. australiensis* (AUS). **(b)** Representative images of different leaf developmental stages of fourth leaves, showing the shoot apical meristem with initiating primordia (SAM+Pi) and successive developmental stages P3, P4, and P5, across the selected four accessions. The bottom panel shows transverse sections of the respective leaf developmental stages used for RNA-sequencing. Scale bars: 1 cm (leaf images) and 100 µm (sections). **(c)** Cleared longitudinal images of leaves showing epidermal cells and stomata at P3, P4, and P5 stages across the four accessions. Scale bar = 20 µm.

**Supplementary Figure 2.**
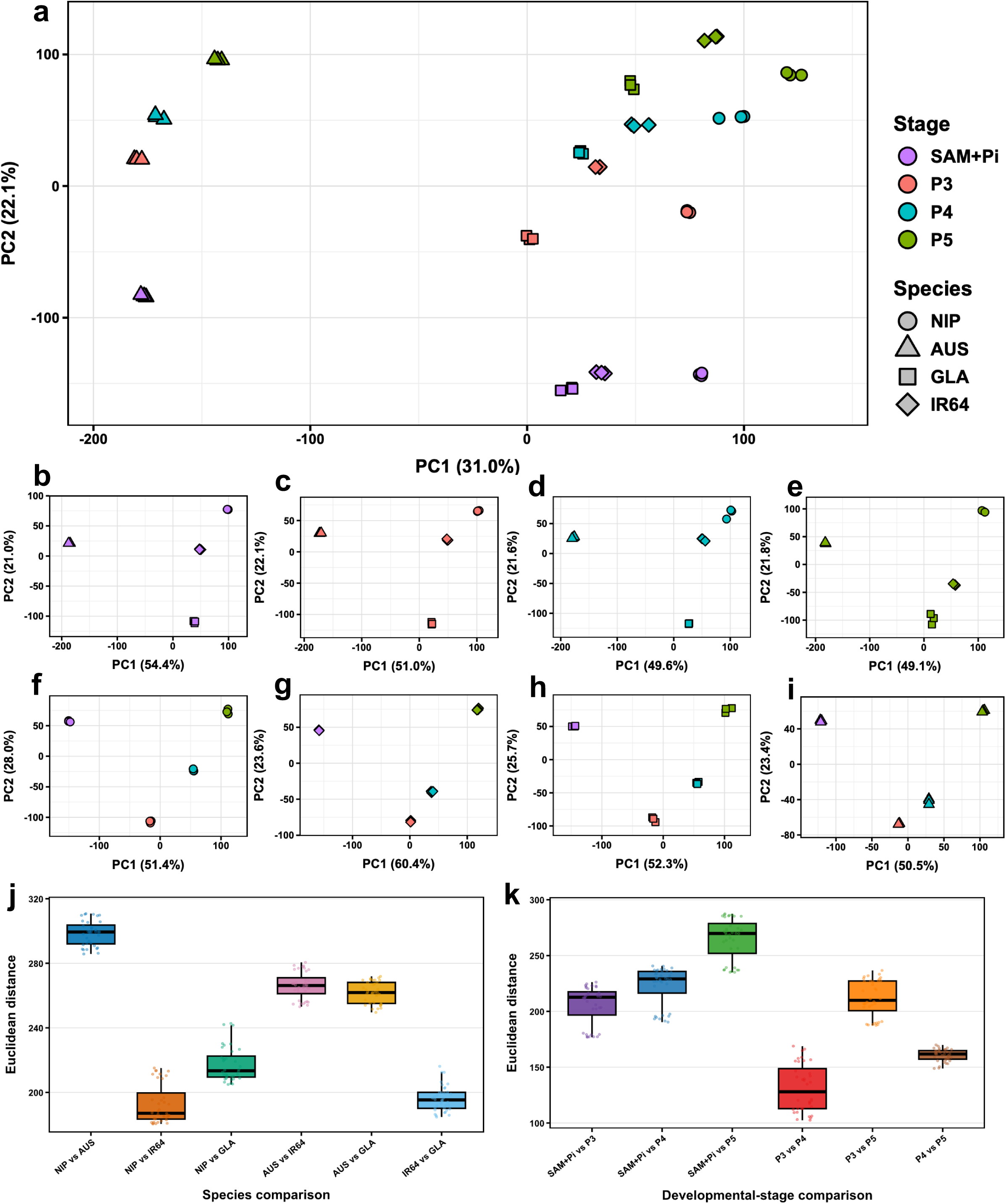
Accession- and stage-specific principal component analysis (PCA) for gene expression profile. **(a)** Principal component analysis (PCA) for all the leaf developmental stages across the selected accessions using normalized gene counts. Each point represents one biological replicate. Colors indicate developmental stages (SAM+Pi, P3, P4, and P5) and shapes indicate accessions. The first two principal components and the percentage of variance explained are shown on the axes. **(b–e)** PCA plots separating different accessions for SAM+Pi **(b)**, P3 **(c)**, P4 **(d)**, and P5 **(e)** developmental stages. **(f–i)** PCA plots separating different developmental stages for *O. sativa* ssp. *japonica* cv. Nipponbare (NIP) **(f)**, *O. sativa* ssp. *indica* cv. IR64 (IR64) **(g)**, *O. glaberrima* (GLA) **(h)**, and *O. australiensis* (AUS) **(i)**. **(j-k)** Euclidean distance calculated between each pairwise species across developmental stages **(j),** Euclidean distance calculated between developmental stage pairs across all species **(k)**.

**Supplementary Figure 3.**
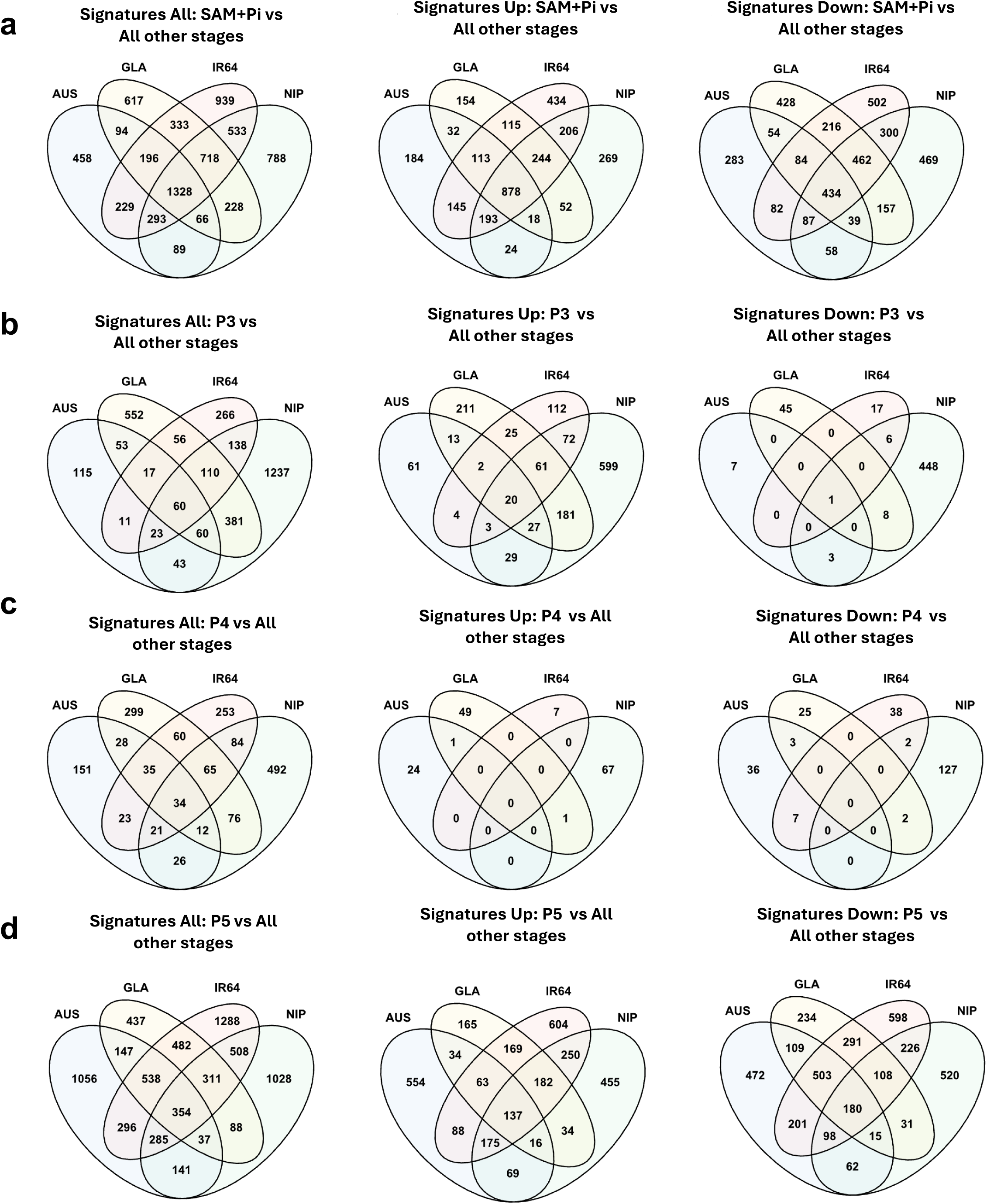
Stage-specific Differentially Expressed Genes (DEGs) across the selected rice accessions. (**a-d**) Venn diagrams showing stage-specific signatures, differentially expressed at a specific developmental stage relative to all other stages across four rice accessions. For each stage - SAM+Pi **(a)**, P3 **(b)**, P4 **(c)**, and P5 **(d)** - three comparisons are shown: all DEGs, upregulated genes, and downregulated genes in the specific developmental stage. Numbers indicate genes shared among accessions.

**Supplementary Figure 4.**
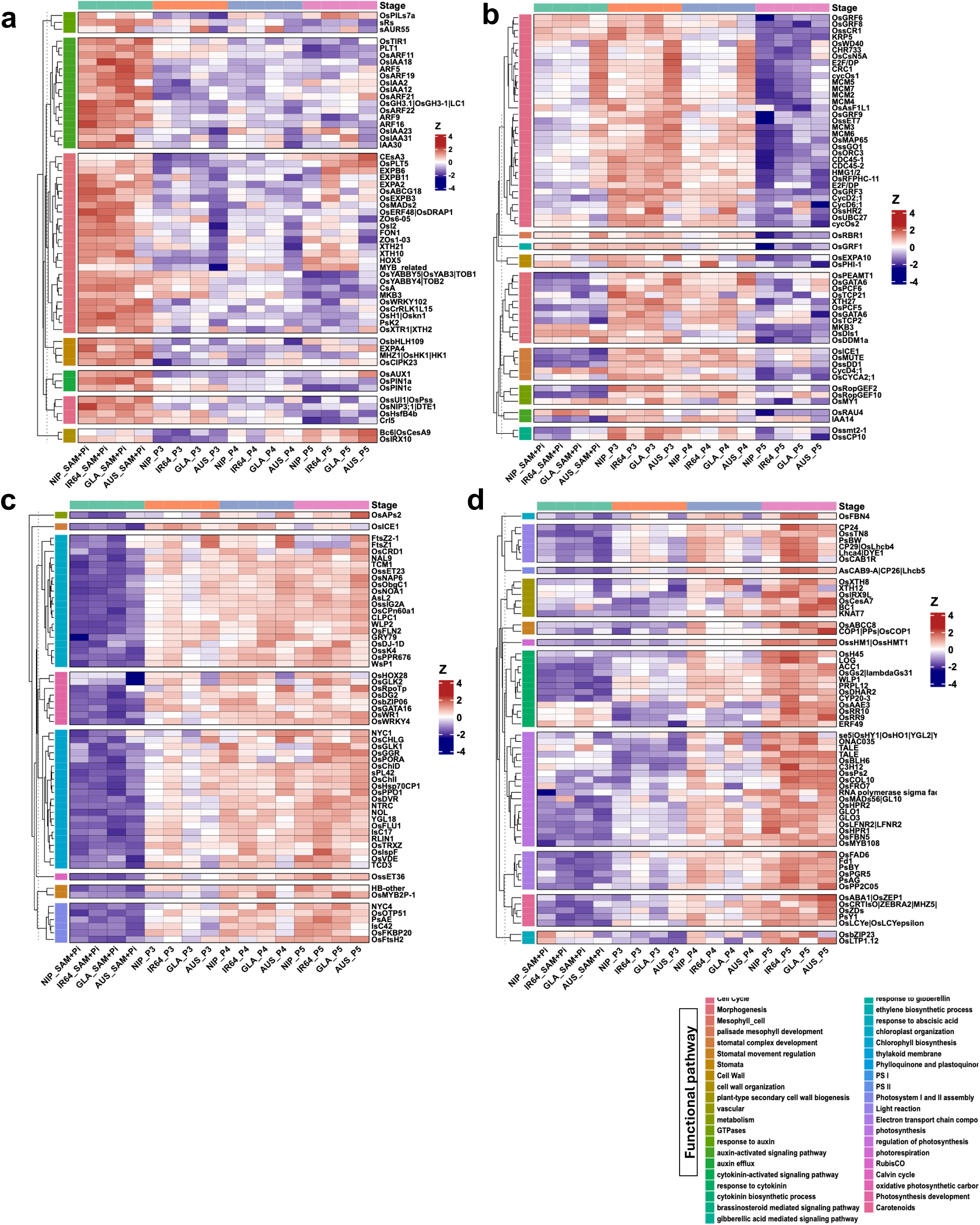
Stage-specific expression patterns of key genes belonging to specific biological pathways across the selected rice accessions. **(a-d)** Heatmaps showing expression patterns of top functional pathway genes predominantly expressed in SAM+Pi **(a)**, P3 **(b)**, P4 **(c)**, and P5 **(d)** developmental stages across the selected rice accessions. Heatmap values correspond to row-wise Z-scores of log₂-transformed normalized count. Genes are grouped by functional pathway categories and indicated by colored annotations.

**Supplementary Figure 5.**
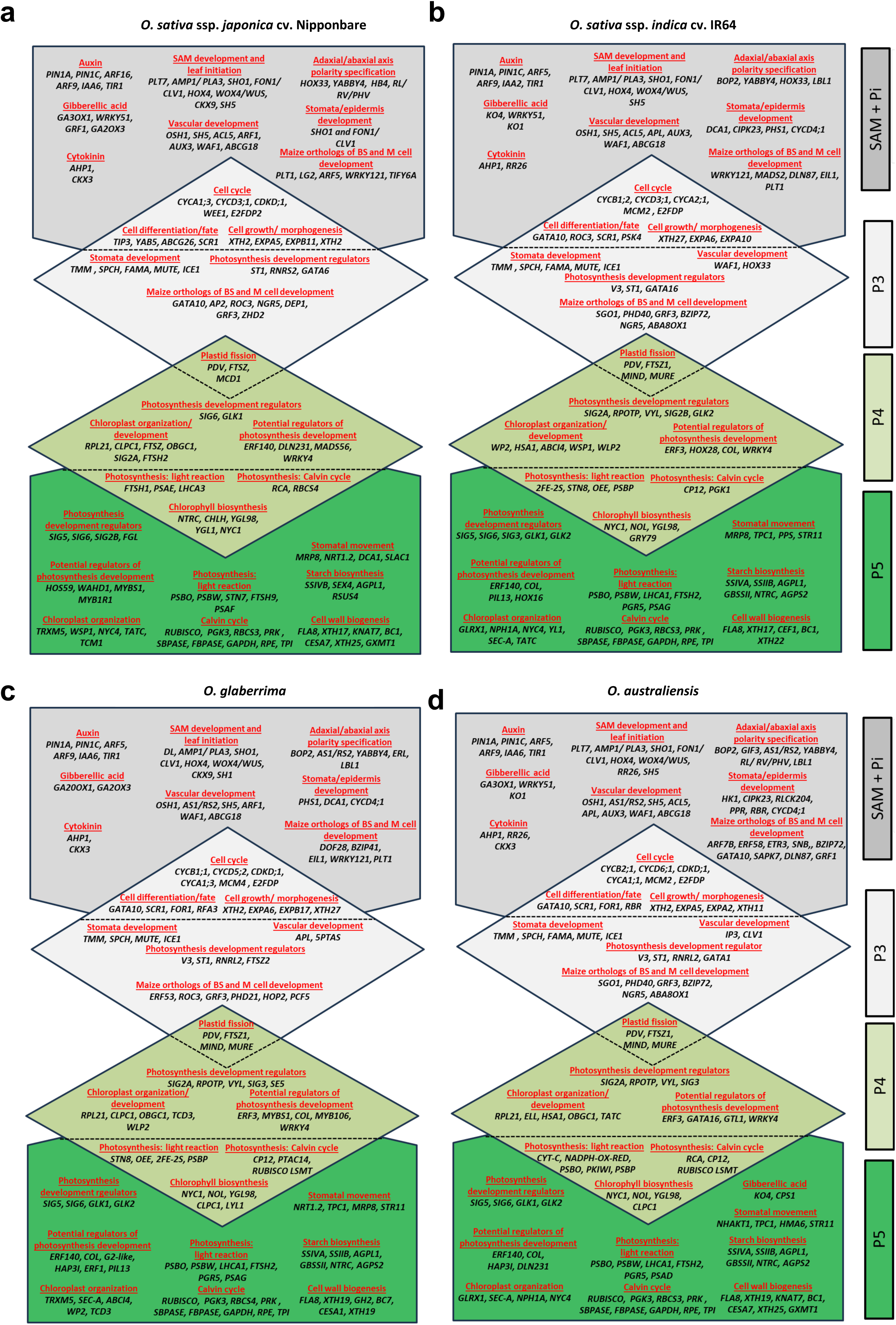
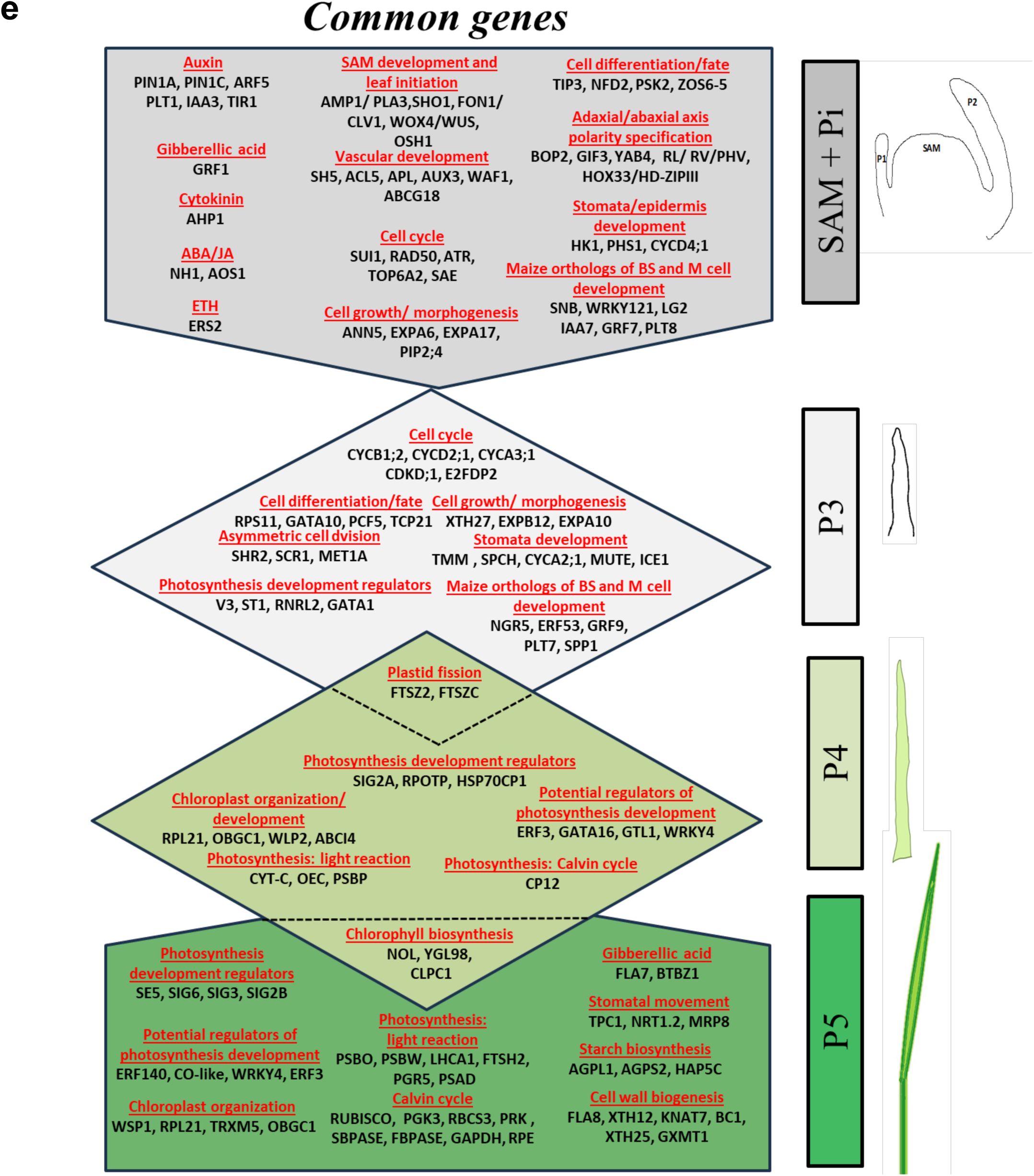
Stage-specific genes and associated functions across the selected rice accessions. The transcriptional gradient summarizes representative stage-specific genes and their putative biological functions across the leaf developmental continuum, from leaf primordium initiation to the establishment of photosynthetic competence. **(a-d)** Accession-specific genes and associated biological functions across the developmental gradients in *Oryza sativa* ssp. *japonica* cv. Nipponbare **(a)**, *O. sativa* ssp. *indica* cv. IR64 **(b)**, *O. glaberrima* **(c)**, and *O. australiensis* **(d). (e)** Genes commonly expressed at specific developmental stages across the selected rice accessions and associated biological functions.

**Supplementary Figure 6.**
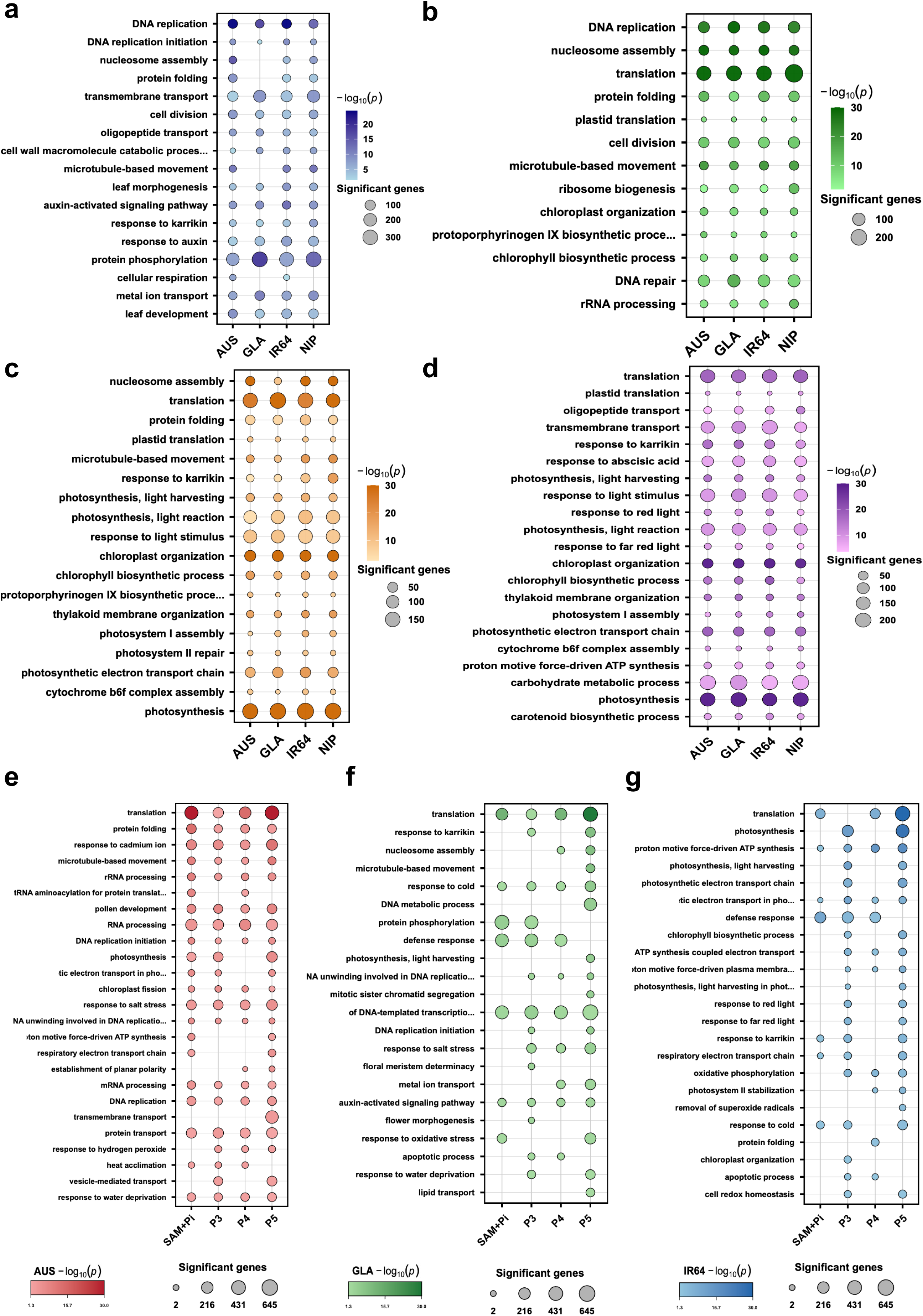
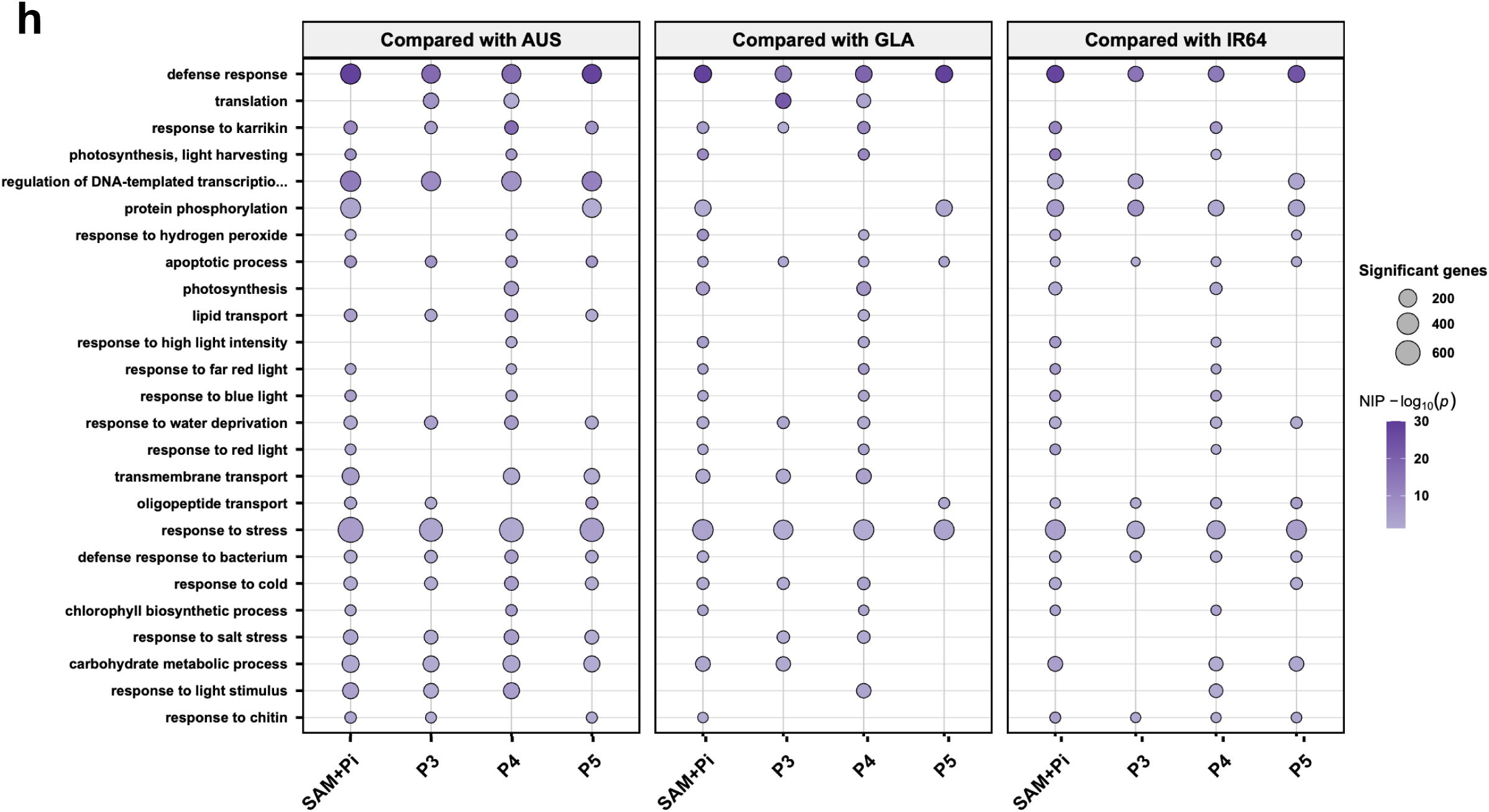
Gene Ontology (GO) enrichment of stage- and accession-specific upregulated genes. **(a–d)** Bubble plots showing GO enrichment for genes upregulated at SAM+Pi **(a)**, P3 **(b)**, P4 **(c)**, and P5 **(d)** across the four rice accessions. **(e–g)** Bubble plots showing GO enrichment for genes upregulated in *O. australiensis* (AUS) **(e)**, *O. glaberrima* (GLA) **(f)**, and *O. sativa* ssp. *indica* cv. IR64 (IR64) **(g)** compared with *O. sativa* ssp. *japonica* cv. Nipponbare (NIP) at each stage. **(h)** GO terms enriched for genes upregulated in NIP compared with the other accessions. Dot size represents the number of genes in each GO term, and color indicates enrichment significance as −log₁₀(P-value).

**Supplementary Figure 7.**
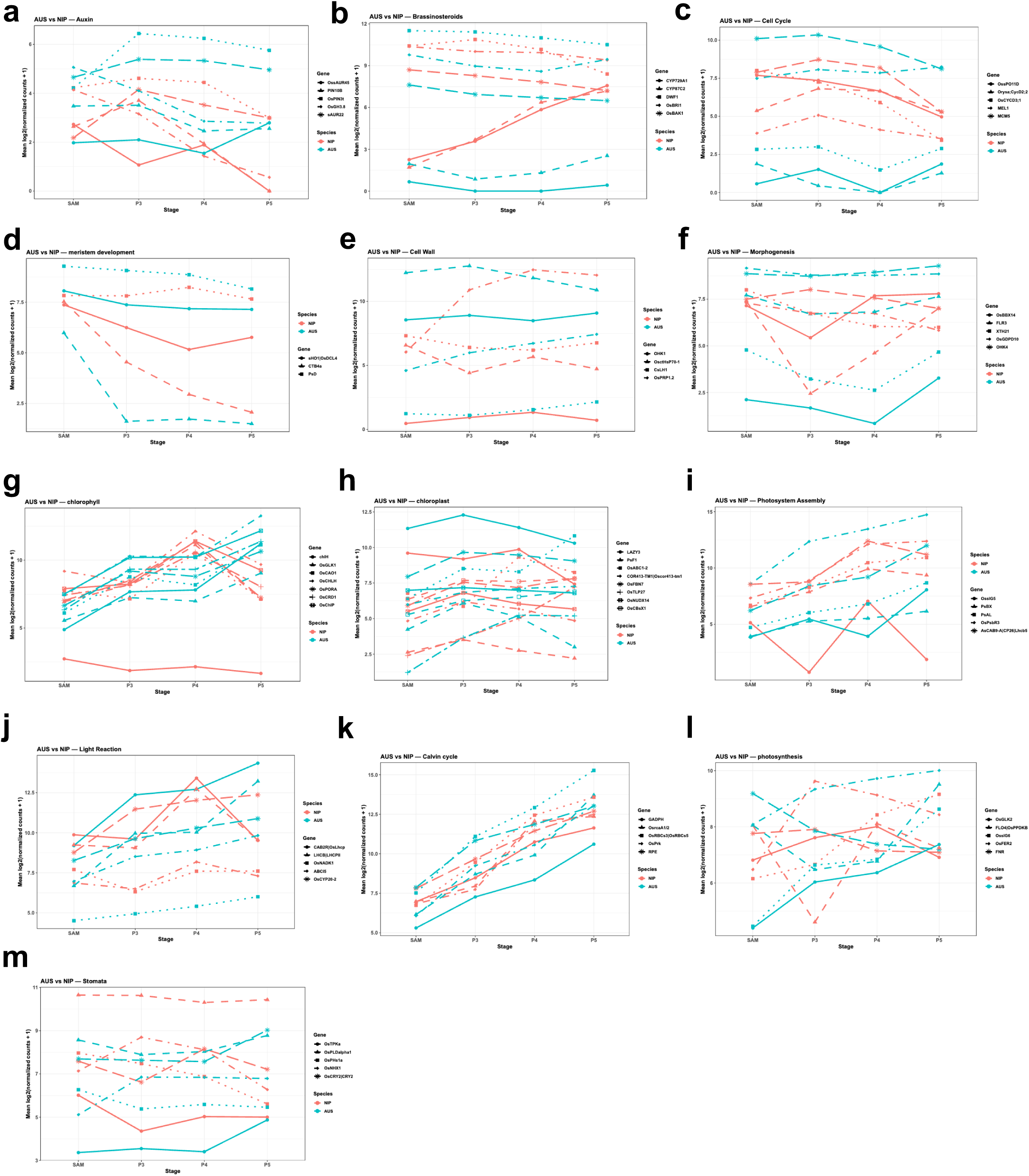
Dynamic expression profile of prominent pathway-related differentially expressed genes (DEGs) across developmental stages in *O. australiensis* (AUS) and *O. sativa* ssp*. japonica* cv. Nipponbare (NIP). **(a-m)** Line plots showing the dynamic expression patterns of the top DEGs associated with major biological pathways across developmental stages in AUS and NIP. Values represent mean log₂-transformed normalized expression levels. Each line corresponds to an individual gene, and colors denote species. Panels represent different biological pathways: Auxin signaling **(a)**, Brassinosteroid signaling **(b)**, Cell cycle **(c)**, Meristem development **(d)**, Cell wall organization **(e)**, Morphogenesis **(f)**, Chlorophyll metabolism **(g)**, Chloroplast organization **(h)**, Photosystem assembly **(i)**, Light reaction **(j)**, Calvin cycle **(k)**, Photosynthesis **(l)**, and Stomatal development and movement **(m)**.

**Supplementary Figure 8.**
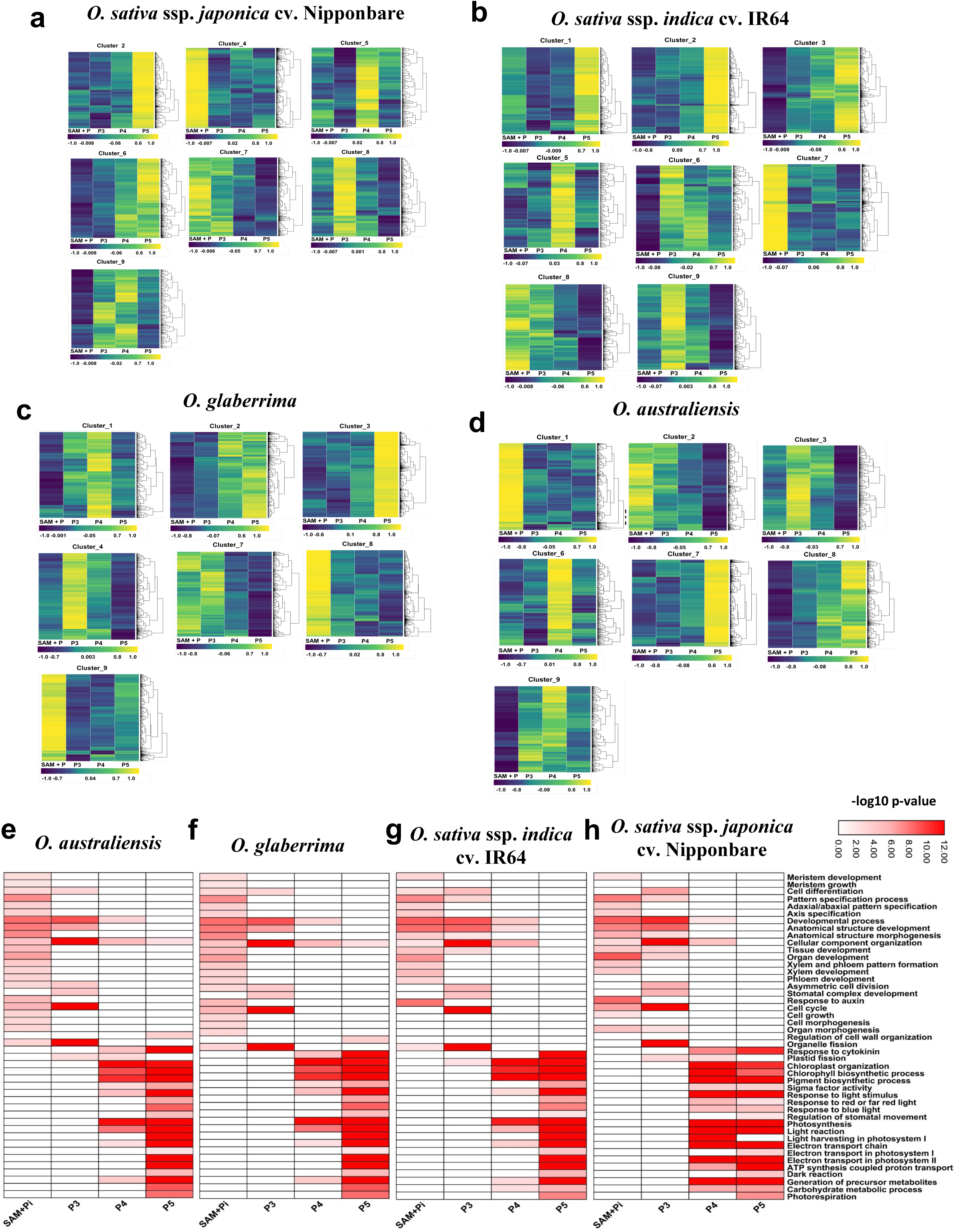
PCA-SOM clustering of gene expression and enriched biological processes at different leaf developmental stages across the selected rice accessions. **(a–d)** Gene expression clusters identified by PCA-SOM analysis across developmental stages for *O. sativa* ssp. *japonica* cv. Nipponbare **(a),** *O. sativa* ssp. *indica* cv. IR64 **(b)**, *O. glaberrima* **(c)**, *O. australiensis* **(d)**. **(e–h)** Enriched biological processes for stage-associated transcript clusters for *O. australiensis* **(e)**, *O. glaberrima* **(f)**, *O. sativa* ssp. *indica* cv. IR64 **(g)**, and *O. sativa* ssp. *japonica* cv. Nipponbare **(h)**.

**Supplementary Figure 9.**
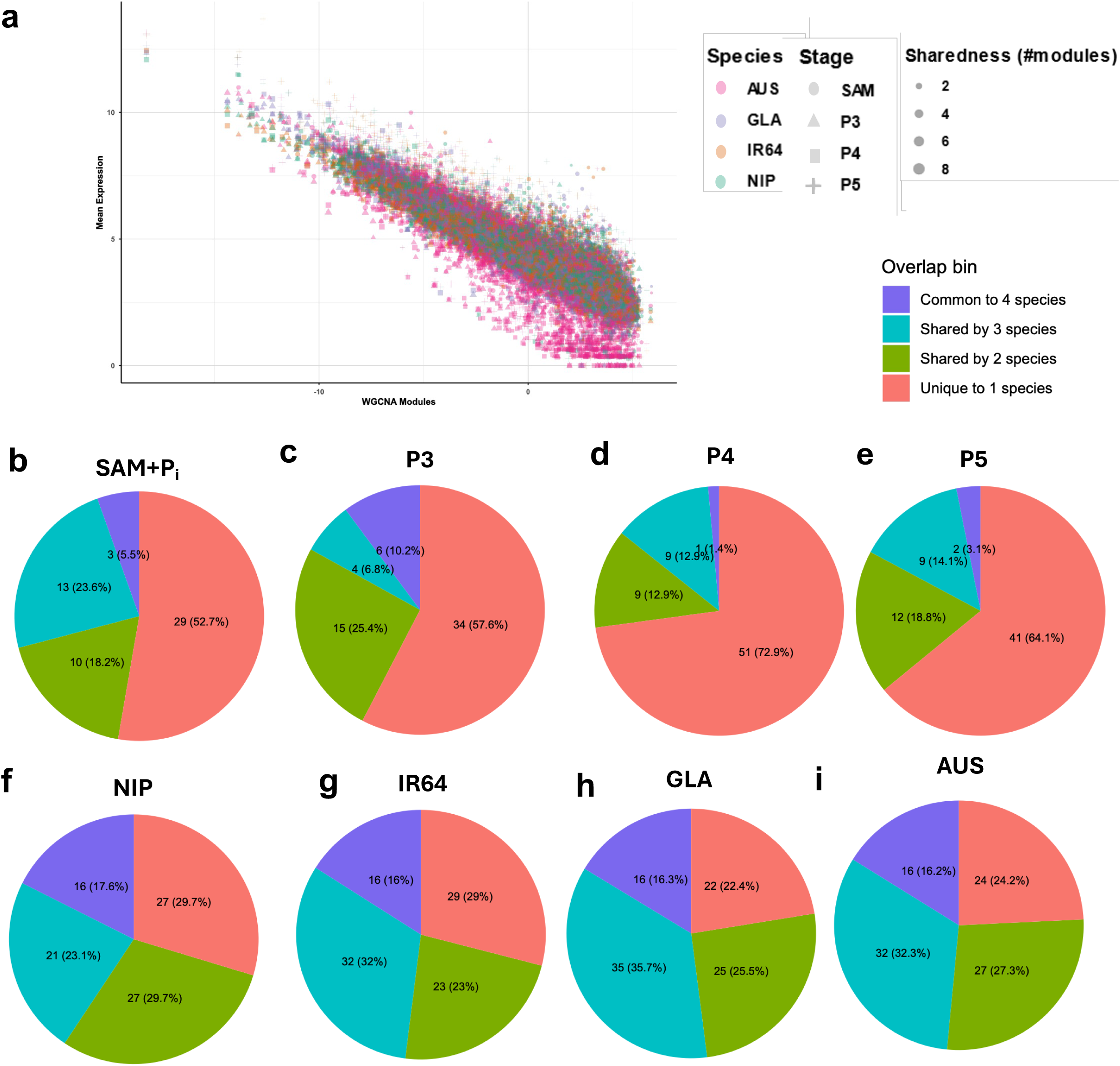
Module-wise gene expression and distribution of hub transcription factors across developmental stages and rice accessions. **(a)** Module-wise gene expression across different developmental stages for the selected rice accessions. The x-axis represents WGCNA-derived co-expression modules, and the y-axis shows mean gene expression. **(b–e)** Overlap of the top 25 hub transcription factors (TFs) across developmental stages: SAM+Pi **(b)**, P3 **(c)**, P4 **(d)**, and P5 **(e)**. **(f–i)** Overlap of the top 25 hub TFs across rice accessions: *O. sativa* ssp*. japonica* cv. Nipponbare (NIP) **(f)**, *O. sativa* ssp. *indica* cv. IR64 **(g)**, *O. glaberrima* **(h)**, and *O. australiensis* **(i)**.

**Supplementary Figure 10.**
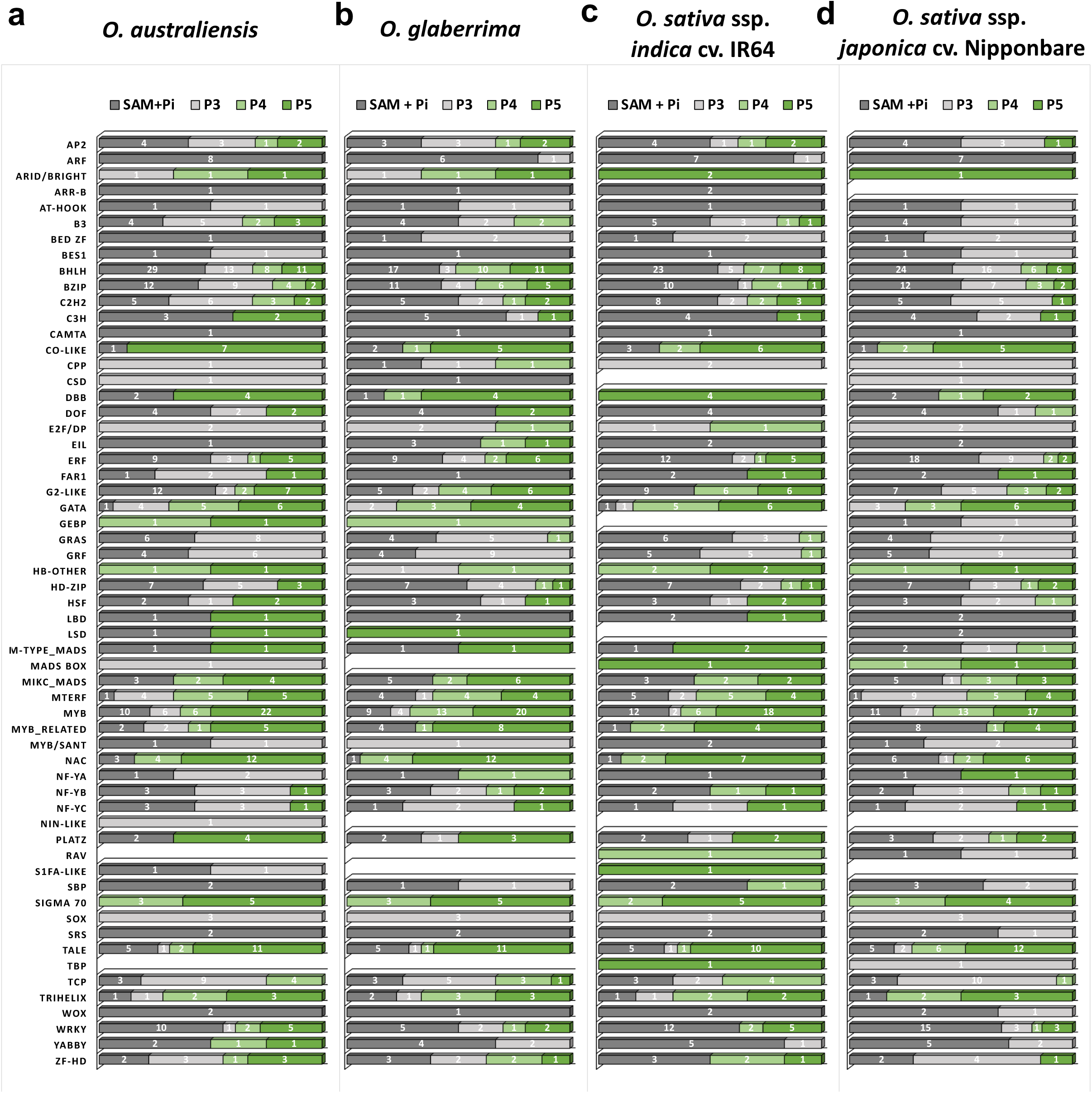
Distribution of transcription factor families across different leaf developmental stages for the selected rice accessions. Distribution of transcription factor (TF) families among hub TFs from WGCNA-derived gene networks at SAM+Pi, P3, P4, and P5 stages in *O. australiensis* **(a)**, *O. glaberrima* **(b)**, *O. sativa* ssp. *indica* cv. IR64 **(c)**, and *O. sativa* ssp. *japonica* cv. Nipponbare **(d)**.

**Supplementary Figure 11.**
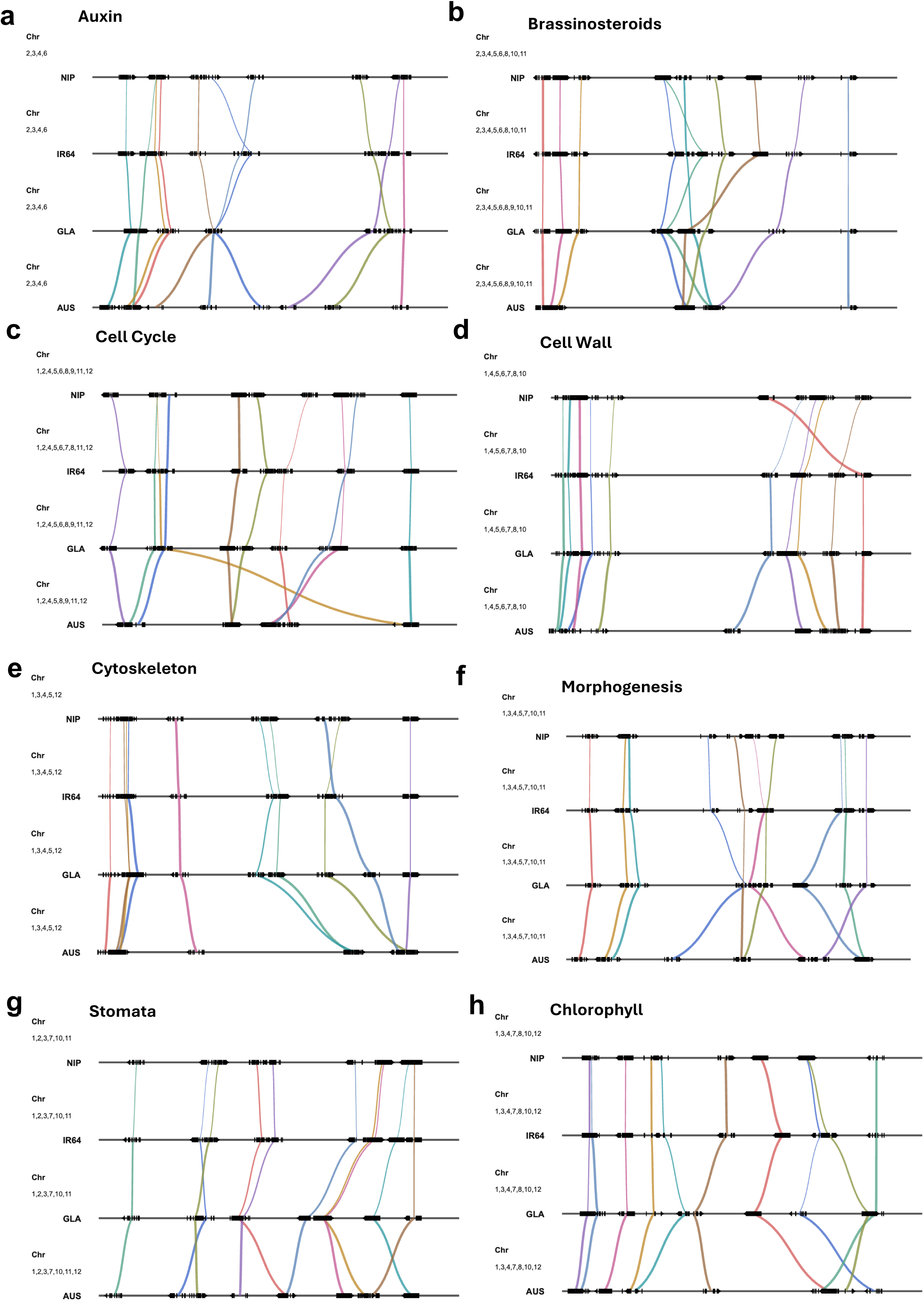
Microsyntenic organization of development- and photosynthesis-associated dynamic DEGs across the four *Oryza* genomes. **(a-h)** Microsyntenic organization of development- and photosynthesis-associated dynamic DEGs across *Oryza sativa* ssp. *japonica* cv. Nipponbare (NIP), *O. sativa* ssp. *indica* cv. IR64 (IR64), *O. glaberrima* (GLA), and *O. australiensis* (AUS). Chromosome numbers are indicated on the left. Horizontal tracks represent genomic intervals, and gene models are displayed with individual blocks representing exons. Coloured links connect collinear gene pairs across genomes, with line thickness representing divergence within the 2-kb upstream regulatory region; thicker lines indicate greater divergence at the promoter. Panels show microsyntenic relationships for genes associated with auxin **(a)**, brassinosteroids **(b)**, cell cycle **(c)**, cell wall **(d)**, cytoskeleton **(e)**, morphogenesis **(f)**, stomatal development **(g)**, and chlorophyll-associated functions **(h)**.

**Supplementary Figure 12.**
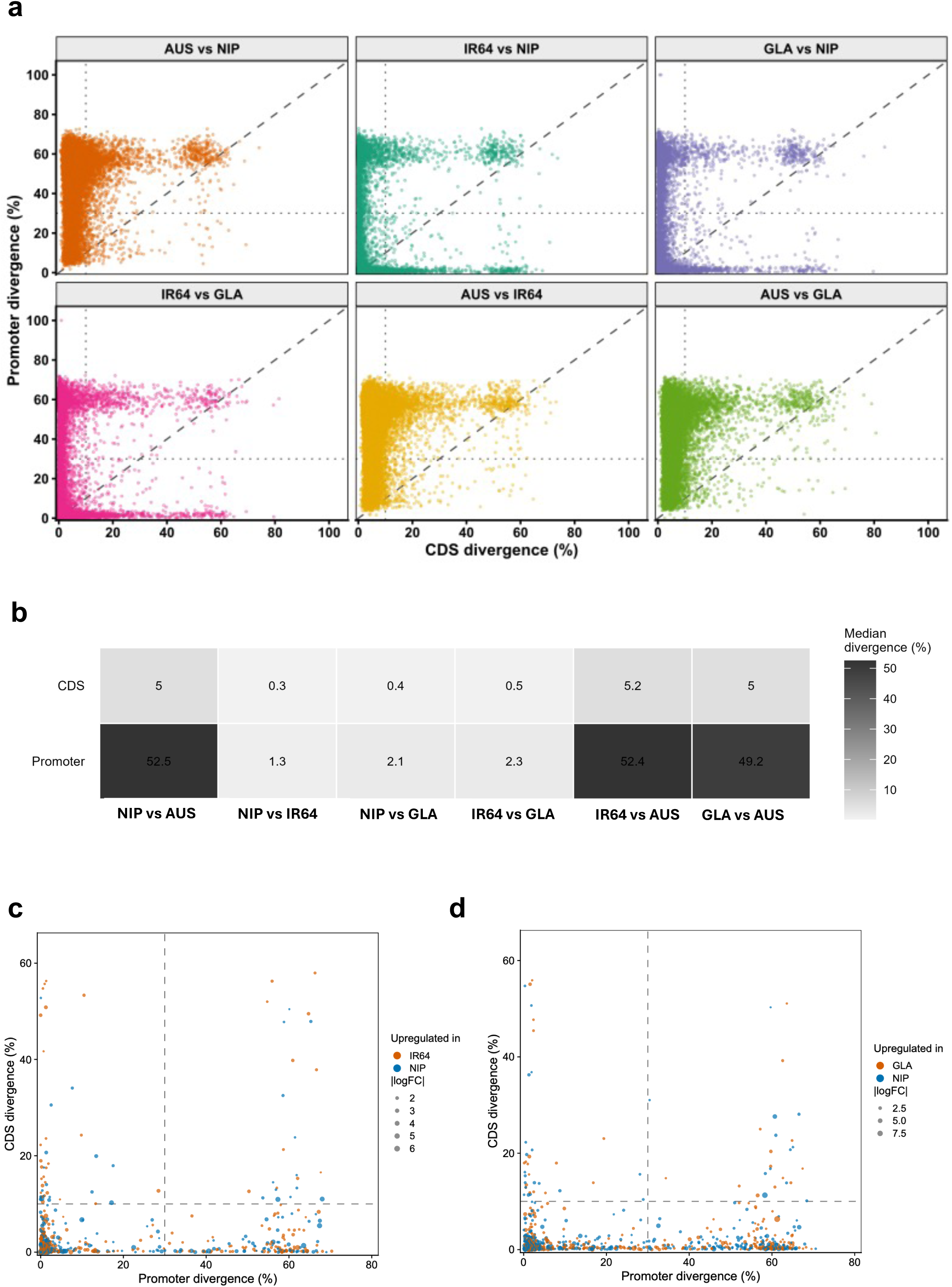
Promoter and coding sequence (CDS) divergence across the selected rice accessions and their association with differential expression. **(a)** Pairwise comparison of promoter and coding DNA sequence (CDS) divergence among *Oryza sativa* ssp. *japonica* cv. Nipponbare (NIP), *O. sativa* ssp. *indica* cv. IR64 (IR64), *O. glaberrima* (GLA), and *O. australiensis* (AUS). Each point represents one orthologous gene pair. The x-axis shows CDS divergence, and the y-axis shows promoter divergence. Dashed lines indicate divergence thresholds used for classification. **(b)** Heatmap showing median promoter and CDS divergence for pairwise comparisons among the accessions. Values indicate median divergence percentage. **(c–d)** Representation of CDS and promoter sequence divergence for differentially expressed genes between NIP and IR64 **(c)** and NIP and GLA **(d)**. Each dot represents one DEG, coloured by the accession in which the gene is upregulated, and the dot size represents absolute log₂ fold change. Dashed lines indicate the thresholds to define low- and high-divergence classes.

**Supplementary Figure 13.**
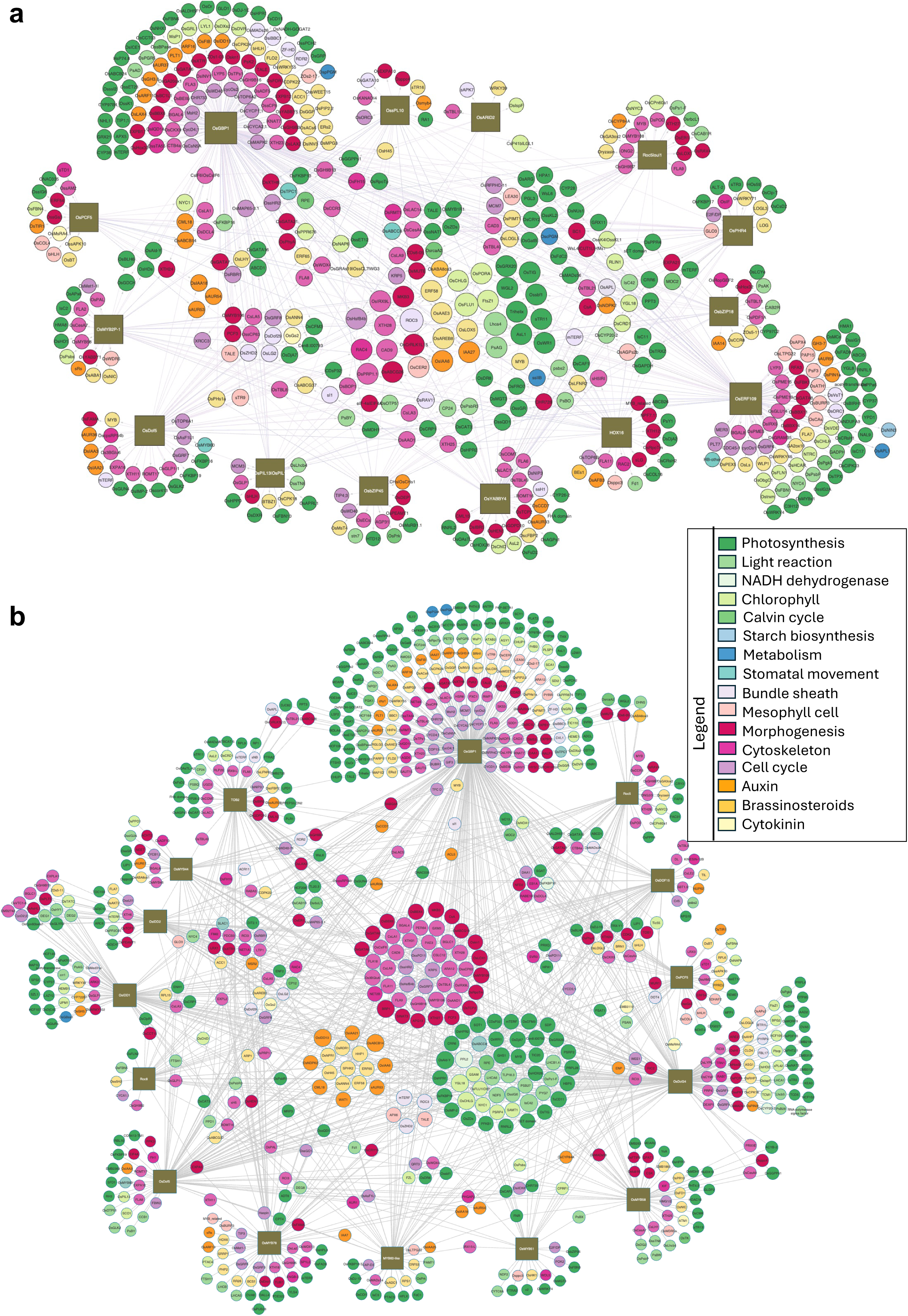
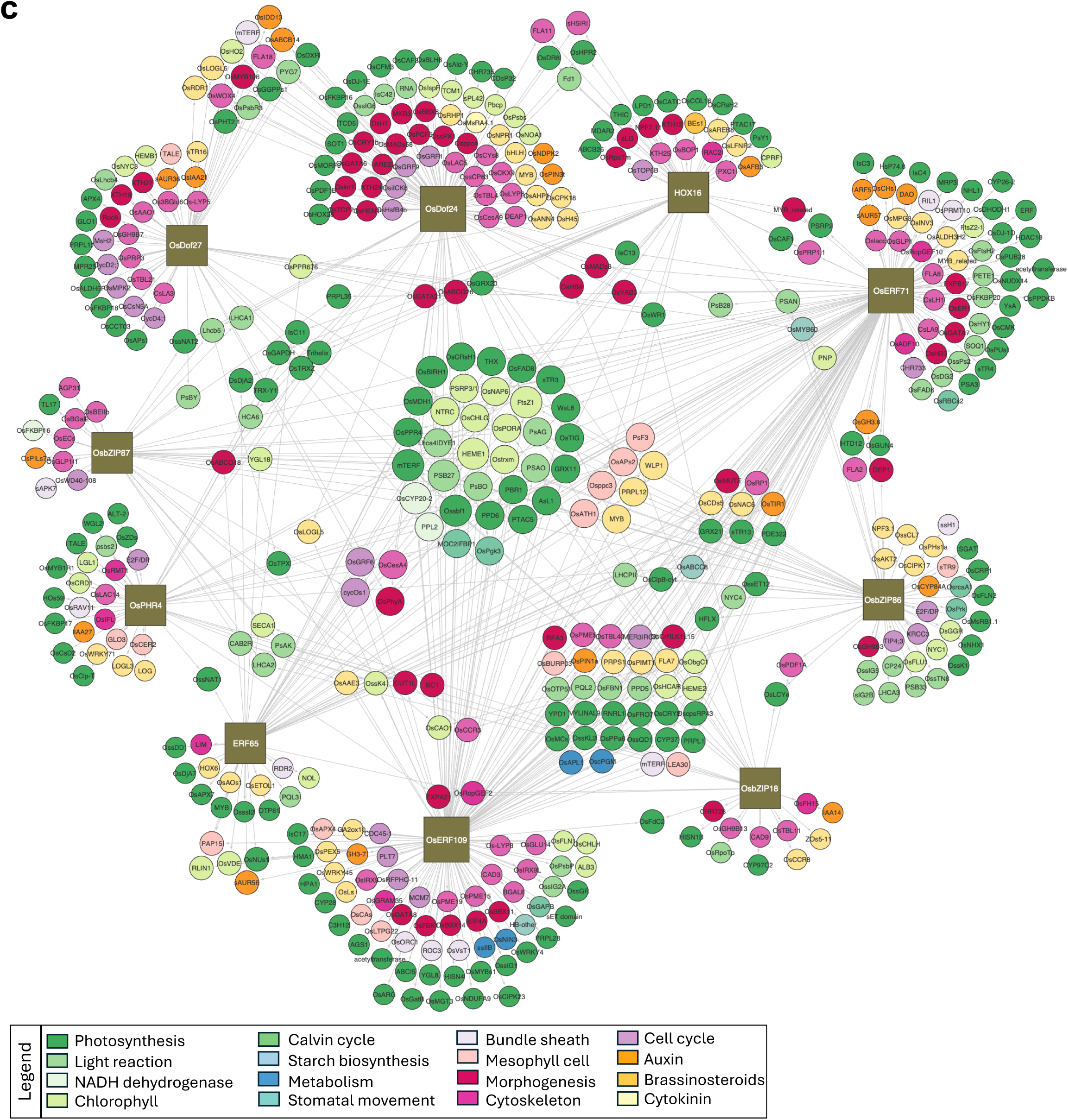
Gene regulatory networks of selected “Category-specific” transcription factors conserved across the four rice accessions. **(a)** Gene regulatory networks of candidate “Category-specific” transcription factors (TFs) and their conserved predicted target genes across the four rice accessions. **(b)** Gene regulatory subnetwork of “developmental” TFs and their predicted conserved target genes across the four rice accessions. **(c)** Gene regulatory subnetwork of “photosynthetic” TFs and their predicted conserved target genes across the four rice accessions. Only regulatory interactions shared across all four accessions are shown. Node colours indicate functional categories of target genes.

**Supplementary Figure 14.**
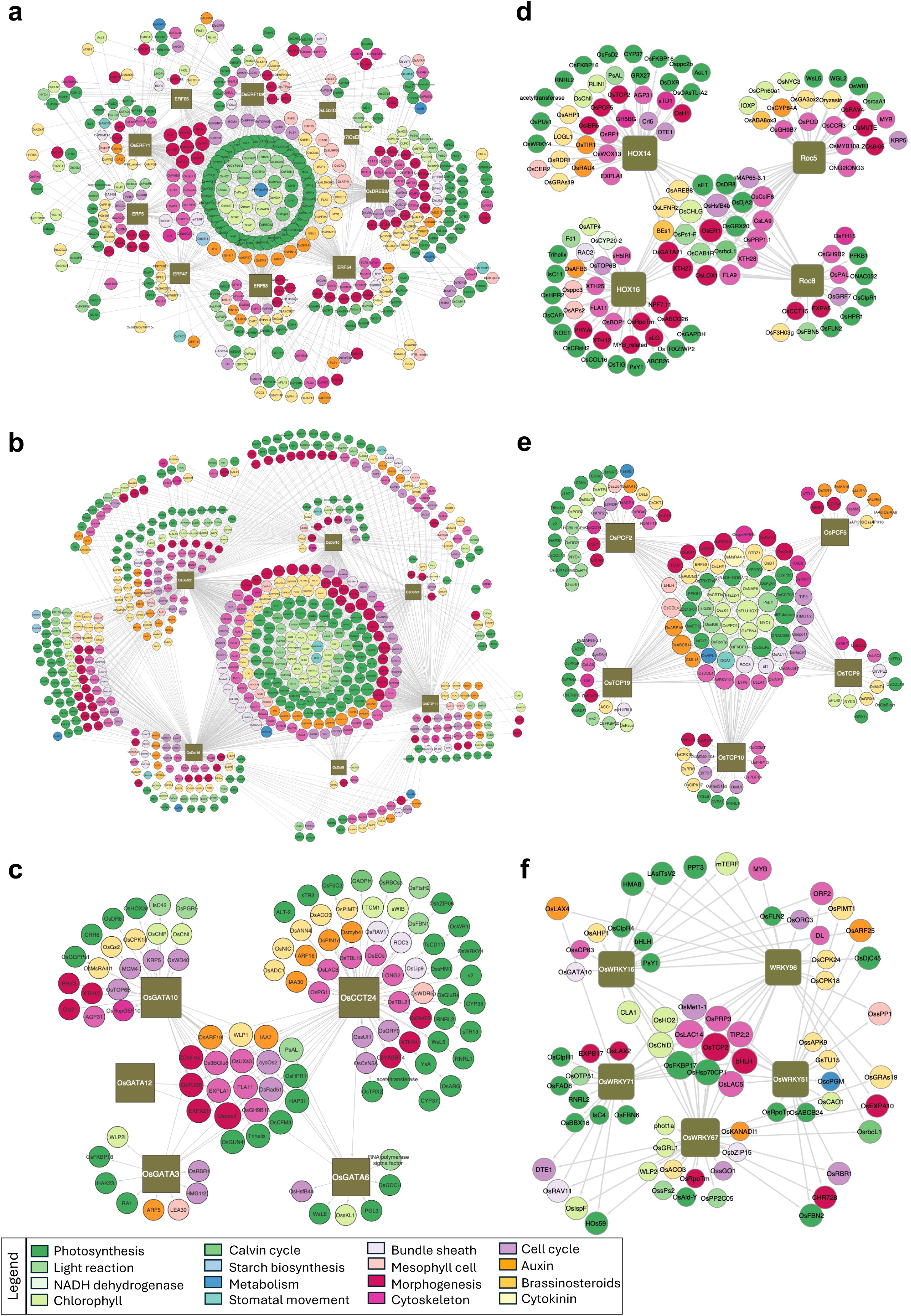
Gene regulatory subnetworks of major transcription factor families and their conserved targets across the four rice accessions. **(a-f)** Gene regulatory subnetworks showing conserved predicted target genes of AP2 **(a)**, DOF **(b)**, GATA **(c)**, Homeodomain **(d)**, TCP **(e)**, and WRKY **(f)** transcription factor families across the four rice accessions. Only regulatory interactions shared across all four rice accessions are shown. Node colours indicate functional categories of target genes.

**Supplementary Figure 15.**
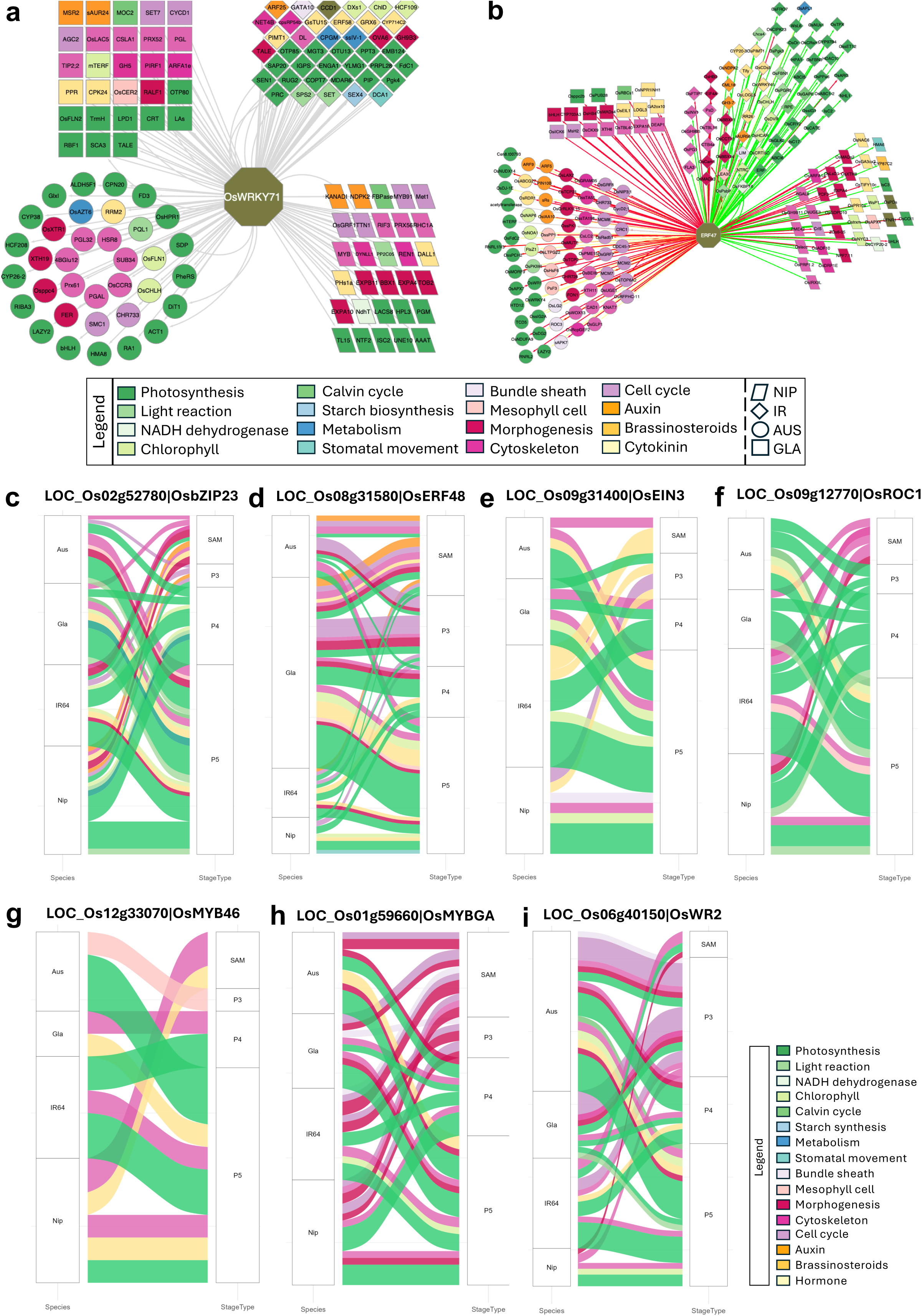
Species-specific regulatory rewiring of key transcription factors. **(a-b)** Gene regulatory subnetworks showing predicted accession-specific target genes of OsWRKY71 **(a)** and ERF47 **(b)**. Node colours indicate functional pathway categories of target genes, and node shapes distinguish accessions. **(c–i)** Alluvial plots showing species-specific predicted target genes of important biological processes across different leaf developmental stages for the transcription factors OsbZIP23 **(c)**, OsERF48 **(d)**, OsEIN3 **(e)**, OsROC1 **(f)**, OsMYB46 **(g)**, OsMYBGA **(h)**, and OsWR2 **(i)**.

**Supplementary Figure 16.**
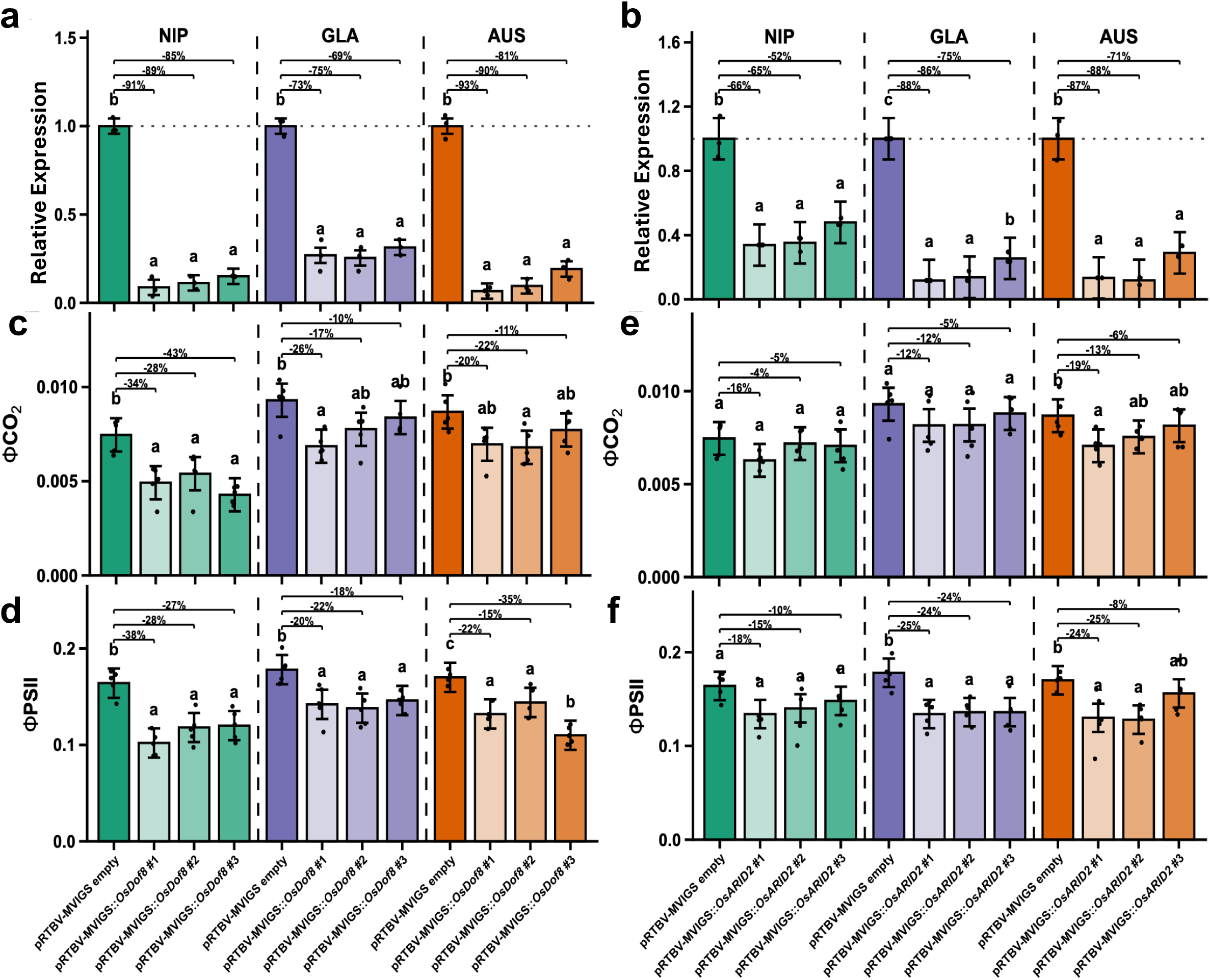
Effects of OsDOF8 and OsARID2 silencing on photosynthetic traits across rice accessions. **(a–b)** Relative transcript expression of *OsDOF8* **(a)** and *OsARID2* **(b)** in *OsDOF8*- and *OsARID2*-silenced NIP, GLA, and AUS plants. **(c-d)** Quantification of ΦCO₂ **(c)** and effective quantum yield of PSII (ΦPSII) **(d)** in *OsDOF8*-silenced *Oryza sativa* ssp. *japonica* cv. Nipponbare (NIP), *O. glaberrima* (GLA), and *O. australiensis* (AUS) plants. **(e-f)** Quantification of ΦCO₂ **(e)** and ΦPSII **(f)** in *OsARID2*-silenced NIP, GLA, and AUS plants. Empty-vector plants were used as controls. Percent values above bars indicate reduction relative to the corresponding empty-vector control. Statistical comparisons were performed using one-way ANOVA followed by Tukey’s HSD test; different letters indicate significant differences at *P* < 0.05. Accessions are color-coded: NIP – green, GLA – purple, and AUS – orange, with shading indicating silencing events.

**Supplementary Figure 17.**
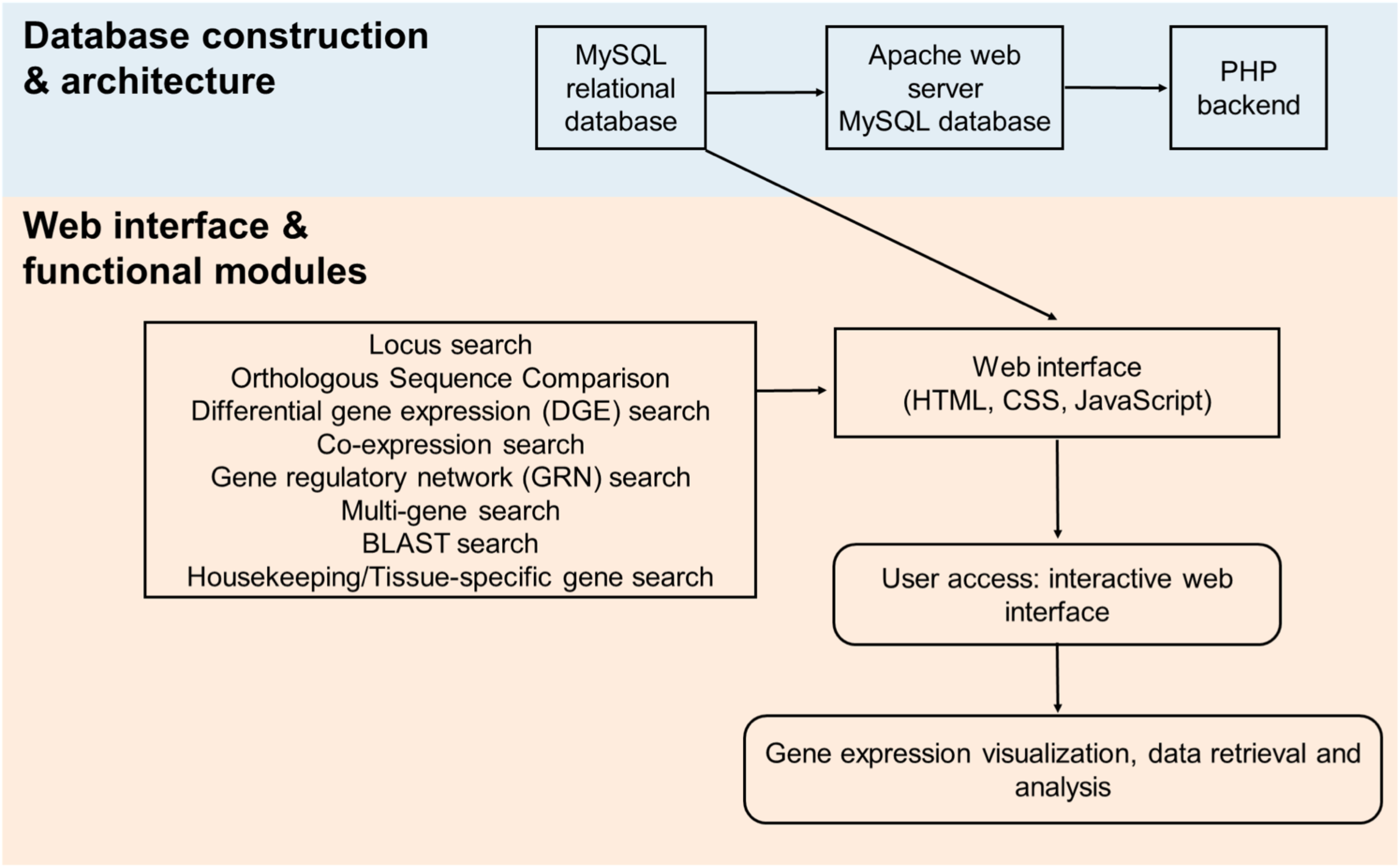
Data integration framework, database architecture, and functional modules of Rice DEV-LEAF.

## Supplementary Tables

**Supplementary Table 1.**
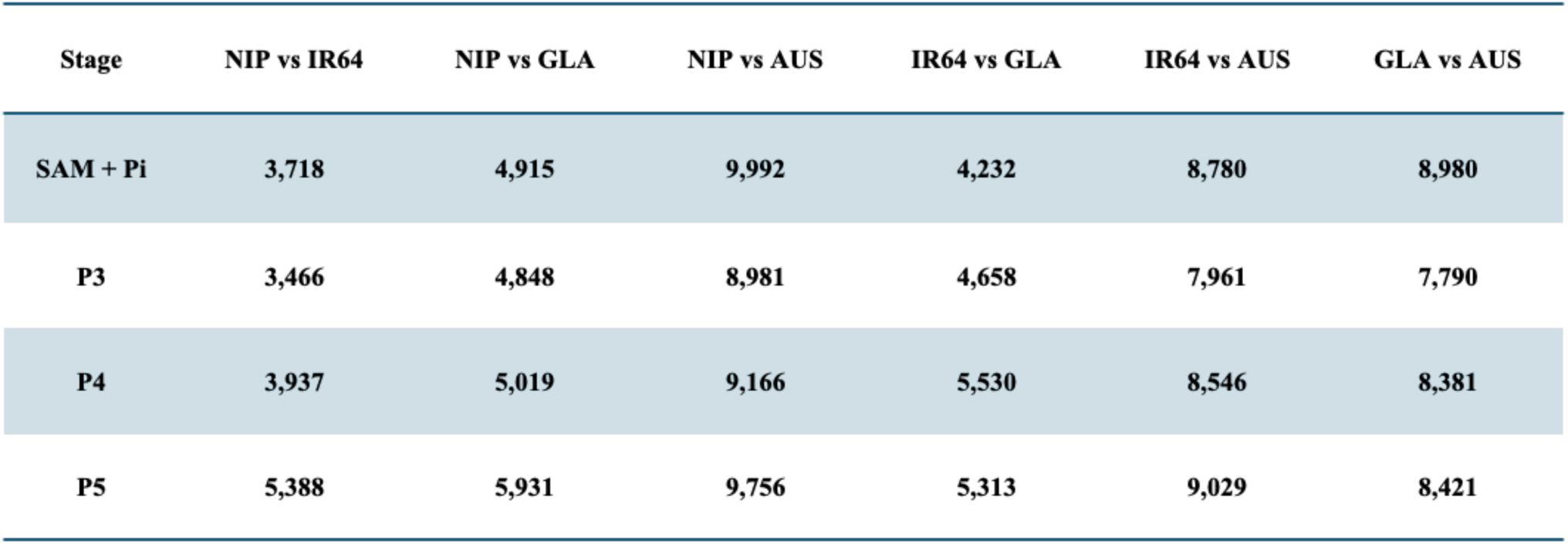
Number of differentially expressed genes between accessions at each developmental stage. Total number of differentially expressed genes identified for each pairwise accession comparison at the SAM + Pi, P3, P4, and P5 developmental stages.

| Stage | NIP vs IR64 | NIP vs GLA | NIP vs AUS | IR64 vs GLA | IR64 vs AUS | GLA vs AUS |
| --- | --- | --- | --- | --- | --- | --- |
| SAM + Pi | 3,718 | 4,915 | 9,992 | 4,232 | 8,780 | 8,980 |
| P3 | 3,466 | 4,848 | 8,981 | 4,658 | 7,961 | 7,790 |
| P4 | 3,937 | 5,019 | 9,166 | 5,530 | 8,546 | 8,381 |
| P5 | 5,388 | 5,931 | 9,756 | 5,313 | 9,029 | 8,421 |

**Supplementary Table 2.** Number of differentially expressed genes between developmental stages within each accession. Total number of differentially expressed genes identified for each pairwise developmental stage comparison in the selected accessions.

| Species | SAM + Pi vs P3 | SAM + Pi vs P4 | SAM + Pi vs P5 | P3 vs P4 | P3 vs P5 | P4 vs P5 |
| --- | --- | --- | --- | --- | --- | --- |
| NIP | 7,832 | 7,403 | 9,454 | 4,419 | 9,140 | 4,484 |
| IR64 | 6,744 | 6,914 | 9,920 | 1,328 | 7,908 | 5,448 |
| GLA | 5,999 | 7,213 | 8,514 | 2,744 | 7,480 | 3,912 |
| AUS | 4,334 | 4,874 | 7,535 | 925 | 6,033 | 3,964 |

## Supplementary Datasets

**Supplementary Dataset S1.** Normalized RNA-seq count matrix across four developmental stages in the four rice species, along with the genome-wide transcriptomic variation represented as a Euclidean distance matrix.

**Supplementary Dataset S2.** List of pairwise differentially expressed genes identified across developmental stages and accessions.

**Supplementary Dataset S3.** Differentially expressed genes for the comparison of leaf developmental stages that are shared across the four rice accessions.

**Supplementary Dataset S4.** Gene Ontology (GO) enrichment results for differentially expressed genes identified in pairwise comparisons between different leaf developmental stages for the selected rice accessions.

**Supplementary Dataset S5.** List of genes and enriched Gene Ontology (GO) categories for each gene coexpression cluster derived from PCA-SOM clustering for the selected rice accessions.

**Supplementary Dataset S6.** List of leaf developmental and photosynthetic genes extracted from PCA-SOM clusters for each leaf developmental stage in the selected rice accessions.

**Supplementary Dataset S7.** List of genes belonging to developmental stage–specific gene co-expression modules identified from WGCNA for the selected rice accessions.

**Supplementary Dataset S8.** Transcription factor (TF) family distribution extracted from gene co-expression networks (GCNs) for each leaf developmental stage across the selected rice accession.

**Supplementary Dataset S9.** Genome-wide collinearity gene-pair dataset across the four rice accessions.

**Supplementary Dataset S10.** Percent identities (PIDs) for coding DNA sequences (CDS) and promoter sequences of high-confidence orthologous genes across the four rice accessions.

**Supplementary Dataset S11.** List of predicted target genes for transcription factors in developmental stage–specific gene regulatory networks (GRNs).

**Supplementary Dataset S12.** List of transcription factors assigned to network topology classes based on the enrichment of pathway categories among their predicted target genes.

**Supplementary Dataset S13.** Gene Ontology (GO) enrichment results for genes predicted as targets of individual transcription factor families.

**Supplementary Dataset S14.** Transcription factors with high cross-species variations in predicted target genes across the selected rice accessions (Cross-Species Target Variation ≥ 0.8).

**Supplementary Dataset S15.** List of primers used in this study.

